# The Patchy Distribution Of The Photosynthetic Gene Cluster In *Erythrobacteraceae*

**DOI:** 10.1101/2025.11.19.689285

**Authors:** Alexandra Grace Walling, Rob DeSalle, Susan L Perkins

## Abstract

The photosynthetic gene cluster which encodes the ability to engage in aerobic anoxygenic photosynthesis is found in multiple clades of *Erythrobacteraceae*, but whether this patchy distribution was caused by horizontal operon transfer, secondary loss, or a mixed regime of transfer and loss has not previously been answered. We use whole genome sequencing and comparative genomic and phylogenetic methods to address these questions. Two new genomes for *Erythrobacteraceae* spp. isolates, *Altererythrobacter* sp. DSM 24483 and *Erythrobacter* sp. DSM 25594, were sequenced, assembled, and annotated. Despite their disparate sampling locations, these genomes are found to be highly similar, differing chiefly in the presence of four genomic islands unique to DSM 24483. A species phylogeny for 91 members of *Erythrobacteraceae,* including the two new genomes, was generated from whole genome data, expanding the most recently published phylogeny for this family by 17 species. Results of the phylogenetic analysis indicate that all species which both possess the photosynthetic gene cluster and express bacteriochlorophyll *a* in culture form one clade, corresponding to the *Erythrobacter* genus. Finally, two hypotheses about the evolution of phototrophy in *Erythrobacteraceae* are tested. Two methods of ancestral state reconstruction and gene order data are leveraged to determine whether secondary loss, horizontal operon transfer, or a mixed regime of transfer and loss explains the patchy distribution of the photosynthetic gene cluster in *Erythrobacteraceae*. Horizontal operon transfer is found to be the best explanation for the evolution of the patchy distribution of the photosynthetic gene cluster in *Erythrobacteraceae*.

## INTRODUCTION

Aerobic anoxygenic photosynthesis is a mode of nutrition in which organic carbon is required for metabolism, but water is not used as an electron donor(Yurkov & Beatty, 1998). Aerobic anoxygenic photosynthetic (AAP) bacteria are obligate heterotrophs that also have a facultative phototrophic mode of metabolism, and as such are important primary producers in marine and freshwater ecosystems, with figures for abundance estimated from 11% of bacteria in the North East Pacific, 0.8-9.4% in the North West Atlantic, 0.8 – 18% in the Mid Atlantic Bight, 0 – 7% in the North Pacific Gyre, 0 – 11.6% in the Baltic Sea, 0.3 – 24.4% in the South Pacific, and 0.1 – 14.8% in the Western Arctic Ocean (Boeuf et al., 2013; Cottrell et al., 2006; Koblížek, 2015; Kolber et al., 2001; Lami et al., 2007; Masin et al., 2006; Sieracki et al., 2006; Yurkov & Hughes, 2017).

AAP is controlled by a photosynthetic gene cluster (PGC) that is 37-42 kilobasepairs in length (Liu et al., 2019; Zheng et al., 2011, 2016). The PGC is widely but patchily distributed among Proteobacteria even within genera, with sister taxa sometimes differing in their capacity for photosynthesis. The extent to which lateral gene transfer or secondary loss has driven the distribution of AAP is unresolved (Imhoff et al., 2018; Lehours et al., 2018). Plasmids bearing the PGC are known in *Roseobacter litoralis*, *Sulfitobacter guttiformis*, *Sulfitobacter noctilucicola*, *Nerelda ignava, Tateyamaria* sp. ANG-S1, and *Oceanicola* sp. HL-35(Brinkmann et al., 2018; Kalhoefer et al., 2011; Petersen et al., 2012). Plasmid-mediated horizontal operon transfer of the PGC has recently been demonstrated in the family *Rhodobacteraceae*, but secondary loss could not be excluded as a contributing factor(Brinkmann et al., 2018; Kloesges et al., 2011). This picture is further complicated by the fact that many PGC in AAP bacteria are not expressed under culture conditions (Fang et al., 2019a; Subhash et al., 2013), and it is not clear whether these PGC have a functional role in these organisms or are a byproduct of pseudogenization or HGT to an incompatible background.

Aerobic anoxygenic photosynthesis was first described in 1979 in what would later be named *Erythrobacter longus (Shiba et al., 1979; Shiba & Simidu, 1982a)*. While AAP is broadly distributed across this group, a minority of documented *Erythrobacteraceae* genomes sequenced to date contain the photosynthetic gene cluster, and the PGC is patchily distributed throughout *Erythrobacteraceae*, with lineages differing in whether they possess the PGC down to the level of the sister species. As such, this group, with substantial prior taxonomic work already completed (Tonon, LAC, Moreira, APB, Thompson, 2014; Xu et al., 2020a), serves as a useful model taxon for understanding the evolutionary dynamics of AAP shaping its patchy distribution.

*Erythrobacteraceae* lineages show cosmopolitan geographic distribution and adaptation to a wide range of marine, freshwater, and terrestrial habitats (**Figure 1**). Species whose genomes were analyzed in this paper were sampled from estuaries in the South China Sea, solar salterns in Odisha, India, sand in the Tengger desert in China, mangrove sediments in Fujian Province, China, a swimming pool in Tokyo prefecture, Japan, deep sea sediments in the Mariana trench, and ice cores from the East Rongbuk Glacier in the Tibetan plateau among other locations (Fang et al., 2019b; Furuhata et al., 2013; Liao et al., 2017; Subhash et al., 2013; Xing et al., 2017; Xu et al., 2020b; Zhao et al., 2017).

**Figure 1:**
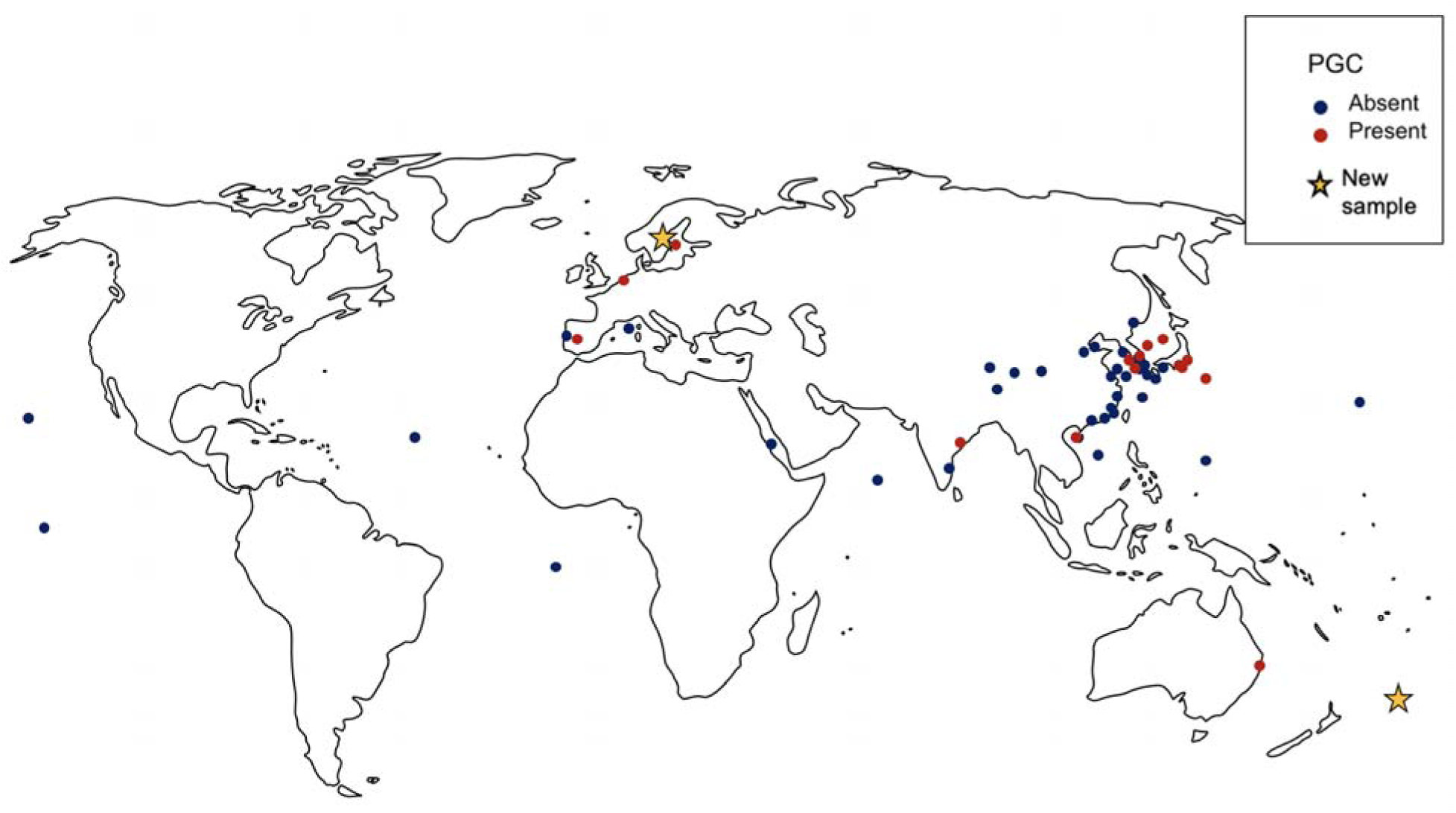
Sampling locations of *Erythrobacteraceae* species whose genomes were studied in this paper.

We use comparative genomic and phylogenetic techniques to explain the patchy distribution of the photosynthetic gene cluster in *Erythrobacteraceae* and illuminate how photosynthesis, an ecologically significant trait, has evolved in this diverse family of bacteria. We sequenced two *Erythrobacteraceae* isolates present in culture collections, *Altererythrobacter* sp. DSM24483 and *Erythrobacter* sp. DSM25594 and assembled and annotated their genomes. Next, we present a phylogenetic analysis of *Erythrobacteraceae*, updating the phylogeny of Xu et al. (2020) to include 17 additional taxa, including the two newly sequenced genomes. A key limitation of the phylogenomic work of Xu et al. was the restriction of genome inclusion only to type species: at the time of publication, 145 *Erythrobacteraceae* genomes were included in the RefSeq database, nearly double the number of genomes included in the study’s phylogeny (O’Leary et al., 2016). As a result, not all the available genetic diversity of *Erythrobacteraceae* was assayed by Xu et al. Both a species phylogeny and a phylogeny for the photosynthetic gene cluster are inferred.

Finally, we use the inferred phylogenies to explain two phenomena: the evolutionary mechanism responsible for the patchy distribution of the photosynthetic gene cluster in *Erythrobacteraceae*, and the restriction of species known to express bacteriochlorophyll *a* in culture to a single clade, the *Erythrobacter* genus. We seek to answer the question, did a scenario of pure secondary loss, of pure horizontal operon transfer, or of a mixed regime of secondary loss and horizontal operon transfer drive the distribution of the photosynthetic gene cluster in *Erythrobacteraceae*? We do not consider a scenario of gain-of-function mutations through convergent evolution because of the complexity of the photosynthetic gene cluster, which is highly conserved across *Alphaproteobacteria* in both gene content and gene order, spans from 35 to 45 genes at 37 to 45 kilobasepairs in length (Zheng et al., 2011, 2016). Both parsimony and maximum likelihood ancestral state reconstruction methods as well as gene order data are leveraged to answer this question. The results of these analyses support the hypothesis that the photosynthetic gene cluster has become patchily distributed because of spread by horizontal operon transfer, in several cases mediated by integrative conjugative elements, and not due to secondary loss or a mixed regime of transfer and loss.

## METHODS

### Cultures and Sequencing

Genomic DNA for two cultures of *Erythrobacteraceae* sp., *Altererythrobacter* sp. DSM 24483 and *Erythrobacter* sp. DSM 25594, was obtained from the Leibniz Institute DSMZ culture collection. *Altererythrobacter* sp. DSM 24483 was initially isolated from the Äspö Hard Rock Laboratory in Äspö, Sweden in 2010. *Erythrobacter* sp. DSM 25594 was isolated from surface seawater in the south pacific gyre and was initially identified as strain SW159(Yin et al., 2013). Paired-end short-read sequence data was obtained from the Illumina MiSeq platform with an average insert size of 250 base pairs for both strains. Read quality was assessed in FastQC (https://www.bioinformatics.babraham.ac.uk/projects/fastqc/) and reads were trimmed with Trimmomatic version 0.39, providing 182x coverage for strain DSM 24483 and 159x coverage for strain DSM 25594(Bolger et al., 2014). Long read data was obtained from the PacBio Sequel I platform with a mean insert length of 6,510 base pairs for strain DSM 24483 and a mean insert length of 5,415 base pairs for strain DSM 25594. Long read data was filtered for sequence length of >1,000 bp, and the bottom 10% of reads were discarded using Filtlong v0.2.1 (https://github.com/rrwick/Filtlong).

### Assembly and Annotation

A mixture of short-read-only, hybrid, and long-read approaches were used to construct assemblies for both *Altererythrobacter* sp. DSM 24483 and *Erythrobacter* sp. DSM 25594. Short-read-only assemblies were constructed using Velvet v1.2.10, ABySS v 2.2.5, and SPAdes v 3.15.3(Bankevich et al., 2012; Simpson et al., 2009; Zerbino & Birney, 2008). Hybrid assemblies were constructed with SPAdes v 3.15.3 using the HybridSPAdes algorithm and Unicycler v0.5.0(Antipov et al., 2016; Wick et al., 2017). Finally, a long-read-first assembly was constructed for *Altererythrobacter* sp. DSM 24483 using Flye v2.8.3 followed by Illumina short read polishing with Polypolish v0.5.0(Kolmogorov et al., 2019; Wick & Holt, 2022). Quality assessment was performed using QUAST v5.1.0 and BUSCO v5.3.2 to determine the highest-quality assembly for both strains(Mikheenko et al., 2018; Simão et al., 2015).

The best assembly for each strain was annotated using the 02-10-2022 build of the Prokaryotic Genome Annotation Pipeline (PGAP) (Tatusova et al., 2016a). Average nucleotide identity between strains DSM24483 and DSM25594 was calculated using alignment free methods with FastANI(Jain et al., 2018). Pairwise genome alignments between DSM24483 and DSM25594 were produced with progressiveMauve(Darling et al., 2010a). Plasmid annotations were manually inspected, and automated predictions were improved using BLAST+ v 2.13.0 and InterPro database to predict functions(Blum et al., 2021; Madden, 2002). Plasmid maps were built using SnapGene v 6.1 (https://www.snapgene.com/).

### Genome Selection and Annotation

For this study, 90 *Erythrobacteraceae* genomes (Table 1) from the RefSeq database of NCBI were selected, with the reference genome for *Citromicrobium bathyomarinum* selected as the outgroup (O’Leary et al., 2016). Genomes were selected both by incorporating those included in the most recent phylogenomic study for *Erythrobacteraceae*, and by downloading the 16S rRNA gene sequences for 145 RefSeq genomes available as of February, 2020; a gene tree was inferred in RAxML to identify Erythrobacteraceae genomes without specific names which were phylogenetically dissimilar to named *Erythrobacteraceae* species for inclusion in this study (Stamatakis, 2014; Xu et al., 2020b). Additionally, two unpublished genomes described in the previous chapter for *Erythrobacter* sp. DSM25594 and *Altererythrobacter* sp. DSM24483 were included. All genomes were annotated or reannotated with the Prokaryotic Genome Annotation Pipeline version released on 02-10-2022 to ensure consistent annotation across the study (Tatusova et al., 2016b).

**Table 1.** Results of *RELAX* hypothesis test that selective pressure has become relaxed on branches where the PGC is not expressed relative to those branches where the PGC is expressed for 30 conserved genes from the photosynthetic gene cluster present in at least 21 out of 23 PGC-bearing taxa.

| Gene | Annotation | Relaxation/intensification parameter K | P-value | Results |
| --- | --- | --- | --- | --- |
| *bchI* | magnesium chelatase ATPase subunit 1 | 0.92 | 0.2544 | No significant evidence for relaxation or intensification of selection |
| *bchD* | **magnesium chelatase subunit D** | **0.84** | **0.0479** | **Relaxation of selection** |
| *bchO* | alpha beta fold hydrolase | 0.95 | 0.4638 | No significant evidence for relaxation or intensification of selection |
| *crtD* | hydroxyneurosporene dehydrogenase | 0.88 | 0.5321 | No significant evidence for relaxation or intensification of selection |
| *crtI* | **phytoene desaturase** | **0.61** | **0.0001** | **Relaxation of selection** |
| *bchC* | chlorophyll synthesis pathway protein | 1.09 | 0.4547 | No significant evidence for relaxation or intensification of selection |
| *bchX* | chlorophyllide a reductase iron protein subunit X | 1.03 | 0.7439 | No significant evidence for relaxation or intensification of selection |
| *bchY* | chlorophyllide a reductase subunit Y | 1.00 | 0.9444 | No significant evidence for relaxation or intensification of selection |
| *bchZ* | **chlorophyllide a reductase subunit Z** | **1.16** | **0.0182** | **Intensification of selection** |
| *pufB* | light harvesting protein | 0.69 | 0.0595 | No significant evidence for relaxation or intensification of selection |
| *pufA* | light harvesting protein | 3.98 | 0.1516 | No significant evidence for relaxation or intensification of selection |
| *pufL* | photosynthetic reaction center subunit L | 1.02 | 0.8069 | No significant evidence for relaxation or intensification of selection |
| *pufM* | **photosynthetic reaction center subunit M** | **0.59** | **0.0000** | **Relaxation of selection** |
| *tspO* | **tryptophan rich sensory protein** | **0.06** | **0.0080** | **Relaxation of selection** |
| *bchP* | geranylgeranyl diphosphate reductase | 1.00 | 1.0000 | No significant evidence for relaxation or intensification of selection |
| *pucC* | **BCD family MFS transporter** | **1.73** | **0.0000** | **Intensification of selection** |
| *chlG* | chlorophyll synthase | 0.84 | 0.1230 | No significant evidence for relaxation or intensification of selection |
| *ppsR* | **transcriptional regulator PpsR** | **0.63** | **0.0000** | **Relaxation of selection** |
| *bchF* | 2 vinyl bacteriochlorophyllide hydratase | 1.00 | 1.0000 | No significant evidence for relaxation or intensification of selection |
| *bchN* | **ferredoxin protochlorophyllide reductase ATP-dependent subunit N** | **1.33** | **0.0034** | **Intensification of selection** |
| *bchB* | ferredoxin protochlorophyllide reductase ATP-dependent subunit B | 1.06 | 0.4132 | No significant evidence for relaxation or intensification of selection |
| *bchH* | magnesium chelatase subunit H | 0.99 | 0.7785 | No significant evidence for relaxation or intensification of selection |
| *bchL* | ferredoxin protochlorophyllide reductase ATP-dependent iron sulfur ATP-binding protein | 0.78 | 0.1519 | No significant evidence for relaxation or intensification of selection |
| *bchM* | magnesium protoporphyrin IX methyltransferase | 1.07 | 0.4943 | No significant evidence for relaxation or intensification of selection |
| *puhA* | photosynthetic reaction center subunit H | 0.64 | 0.6660 | No significant evidence for relaxation or intensification of selection |
| *puhB* | **PH domain containing protein** | **2.60** | **0.0001** | **Intensification of selection** |
| *puhC* | photosynthetic complex assembly protein | 1.90 | 0.0981 | No significant evidence for relaxation or intensification of selection |
| *acsF* | **magnesium protoporphyrin IX monomethyl ester oxidative cyclase** | **1.25** | **0.0009** | **Intensification of selection** |
| *puhE* | DUF3623 domain containing protein | 0.93 | 0.3047 | No significant evidence for relaxation or intensification of selection |

### Pan-Genome Analysis

Pan-genome prediction, ortholog identification, and multiple sequence alignment for downstream gene tree inference was performed using the PanX pipeline (Ding et al., 2018a). The following parameters were used: DIAMOND was used for clustering with a default e-value threshold of 0.001 and a strict core genome threshold of 1.0. Multiple sequence alignments of amino acid sequences for 120 single copy orthologs identified by PanX present in all 91 genomes were downloaded. Additionally, copies of the 5S, 16S, and 23S rRNA gene for all 91 genomes were extracted from GenBank files generated during annotation and aligned using MUSCLE (Edgar, 2004).

### Phylogenomic Inference

Individual gene trees for all 120 single copy orthologs were inferred using RAxML with the GAMMA model of rate heterogeneity, a maximum likelihood estimate of base frequencies, and the LG substitution model (-m PROTGAMMALGF). Gene trees for the 5S, 16S, and 23S rRNA genes were inferred using the GAMMA model of rate heterogeneity and the GTR substitution matrix (-m GTRGAMMA). 100 bootstrap replicates were computed for all gene trees. A species tree was inferred from the 123 gene trees constructed above using Astral-III (Zhang et al., 2018). Low support branches with support values of 10% or less on gene trees were contracted prior to computing the summary tree with ASTRAL-III. Branches were associated with metadata (presence or absence of the photosynthetic gene cluster in the genome) in the ggtree suite of tools in the R language (Yu et al., 2017).

A phylogeny for the photosynthetic gene cluster was inferred using the methods described above for 25 genes across 19 taxa (Supplementary Table 2). Alignments for amino acid sequences were produced in PanX, individual gene trees were inferred in RAxML, and a consensus tree was computed using ASTRAL-III.

### Ancestral State Reconstruction

Ancestral state reconstruction for the presence or absence of the photosynthetic gene cluster in *Erythrobacteraceae* was performed using the Mesquite software (Maddison & Maddison, 2021). Both parsimony and maximum likelihood methods were used to trace the character history of this binary character data onto the species tree. For parsimony, gains and losses of the photosynthetic gene cluster were weighted equally. For maximum likelihood, the Markov k-state 1 parameter model was used, with both gains and losses weighted equally.

### Multiple Genome Alignment of Photosynthetic Gene Cluster and Flanking Regions

To assess potential transitions between presence or absence of phototrophy among those branches where ancestral state reconstruction returned multiple equally parsimonious scenarios, we compared the genomic regions flanking the photosynthetic gene cluster across taxa using pairwise and multiple genome alignments using progressiveMauve in order to assess whether PGC flanking regions were conserved across ambiguously reconstructed clades (Darling et al., 2010b). Gene order data for the PGC and flanking regions was extracted from GFF files produced during annotation. Automated annotations were manually improved through BLAST+ searches (Camacho et al., 2009). The locations of coding sequences and tRNA genes were extracted using a custom BASH script. Gene order data was plotted using the gggenes R package (Wilkins & Kubota, 2022).

### Test for Relaxation of Selection

To test the hypothesis that selective pressure on the photosynthetic gene cluster has become relaxed on branches where the PGC is not expressed relative to those branches where bacteriochlorophyll *a* has been documented as expressed in culture, I used the RELAX algorithm as implemented in the Hyphy framework for evolutionary hypothesis testing (Kosakovsky Pond et al., 2020; Wertheim et al., 2015). A dataset of 30 genes from the photosynthetic gene cluster and present in at least 21 of 23 taxa was used (**Supplmentary Table 3**). Single-gene alignments were produced in MAFFT through the PanX pipeline as described above (Ding et al., 2018b; Katoh et al., 2002). Alignments were concatenated in Mesquite, producing a DNA supermatrix of 36923 positions, of which 23 positions were removed for representing gaps in the alignment across all 23 taxa (Maddison & Maddison, 2021). A tree of maximum likelihood was inferred for 23 taxa using RAxML, with the GAMMA model of rate heterogeneity and the GTR substitution matrix (-m GTRGAMMA), with 100 bootstrap replicates (Stamatakis, 2014). For RELAX, 12 branches, corresponding to the PGC-expressing organisms in this study, were designated as the reference set (*Erythrobacter* sp. NAP1, *Erythrobacter longus, Erythrobacter dokdonensis, Erythrobacter tepidarius, Erythrobacter sanguineus, Erythrobacter ramosus, Erythrobacter colymbi, Erythrobacter donghaensis, Erythrobacter neustonensis*, *Porphyrobacter* sp. LM 6, *Erythrobacter cryptus, Erythrobacter litoralis* DSM8509), 10 branches were designated as the test set (*Alteriqipengyuania lutimaris, Aurantiacibacter spongiae, Pontixanthobacter sediminis, Pontixanthobacter aestiaquae, Aurantiacibacter marinus, Aurantiacibacter odishensis, Aurantiacibacter zhengii, Altererythrobacter* sp. BO 6, *Altererythrobacter ishigakiensis, Altererythrobacter insulae*), and the outgroup *Citromicrobium bathyomarinum* as well as all internal nodes were designated the unclassified (nuisance) set.

To examine heterogeneity in selective pressure across the photosynthetic gene cluster, the RELAX algorithm was also run for all 30 genes individually. Gene trees were inferred from individual alignments in RAxML, with the GAMMA model of rate heterogeneity and the GTR substitution matrix (-m GTRGAMMA), with 100 bootstrap replicates. The same sets of branches as in the concatenated analysis were designated as reference, test, and unclassified sets.

## RESULTS

### *Assembly of* Altererythrobacter *sp. DSM24483* and Erythrobacter *sp. DSM25594*

Best assemblies for both *Altererythrobacter* sp. DSM24483 and *Erythrobacter* sp. DSM25594 were chosen by selecting the assembly with the smallest number of contigs and the highest N50 for each genome. For *Altererythrobacter* sp. DSM24483, the long-read assembler Flye followed by Illumina short-read polishing with Polypolish achieved the best quality statistics, with eight contigs and an N50 of 3,429,213, for a total length of 3,668,998 base pairs. The GC% was calculated to be 60.84%. Genome completeness was assessed in BUSCO, finding 122/124 complete BUSCOs and 2/124 fragmented BUSCOs, with no missing single copy orthologs. For *Erythrobacter* sp. DSM25594, the hybrid Unicycler assembler achieved a complete genome, with four circular contigs representing a circular chromosome and three plasmids, an N50 of 3,428,904, and a total length of 3,551,136 base pairs. The GC% was calculated to be 60.79%. BUSCO genome completeness assessment found 122/124 complete BUSCOs, 2/124 fragmented BUSCOs, and no missing single copy orthologs.

### *Annotation of* Altererythrobacter *sp. DSM24483* and Erythrobacter *sp. DSM25594*

Functional annotation for *Altererythrobacter* sp. DSM24483 revealed a total of 3,629 coding sequences, 47 tRNA genes, and one copy each of the 5S, 16S, and 23S rRNA genes. The 16S rRNA gene sequence of strain DSM24483 has 97.83 percent identity with that of *Aurantiacibacter zhengii* strain V18 (synonymous with *Erythrobacter zhengii*), isolated from deap sea sediment (GenBank accession number NR_171422.1) according to a BLASTn search using the 16S rRNA gene sequence of strain DSM24483 as a query and using strain V18 as a reference(Fang et al., 2019c). According to the annotation, *Altererythrobacter* sp. DSM24483 lacks the genes encoding the photosynthetic gene cluster responsible for the aerobic anoxygenic photosynthesis phenotype.

Functional annotation for *Erythrobacter* sp. DSM25594 revealed a total of 3,507 coding sequences, 46 tRNA genes, and one copy each of the 6S, 16S, and 23S rRNA genes. The 16S rRNA gene sequence of strain DSM25594 has 99.46 percent identity with that of *Aurantiacibacter zhengii* strain V18 according to a BLASTn search using the 16S rRNA gene sequence of strain DSM25594 as a query and using strain V18 as a reference. According to the annotation, *Erythrobacter* sp. DSM25594 lacks the genes encoding the photosynthetic gene cluster responsible for the aerobic anoxygenic photosynthesis phenotype.

A BLASTn pairwise alignment of the 16S rRNA gene for *Altererythrobacter* sp. DSM24483 and *Erythrobacter* sp. 25594 shows the gene differs in 31 out of 1487 positions, with a percent identity of 97.92. Average nucleotide identity between DSM25594 and DSM24483 was calculated to be >99.99% by FastANI. Across 120 single copy orthologs, all positions in all amino acid sequence alignments were identical. Pairwise genome alignment reveals that strain DSM24483 possesses four genomic islands not found on DSM25594 of lengths 47,514, 20,952, 44,296, and 3,770 base pairs, and a major inversion 715,012 base pairs long on the main chromosome.

### Pan Genome Analysis

The PanX pipeline predicts 37,013 total genes in the *Erythrobacteraceae* pangenome, of which 383 form the core genome present in all 90 *Erythrobacteraceae* sp. and *Citromicrobium bathyomarinum*. Of the 383 core genes, 120 exist as single copy orthologs while 263 are duplicated. The majority of genes in the accessory genome are present in only one genome included in the phylogenetic dataset.

### Phylogenomic Inference

Phylogenomic inference through consensus methods produced a species tree (**Figure 2**) of 91 taxa from 123 single copy orthologs. Mapping associated metadata of presence or absence of the photosynthetic gene cluster from the relevant genome reveals the scattered distribution of the photosynthetic gene cluster across eight clades. Those species which both possess the photosynthetic gene cluster and express bacteriochlorophyll *a* form a monophyletic group. This monophyletic group is recapitulated in the phylogeny for the photosynthetic gene cluster itself (**Figure 3**), albeit with some rearrangements in the position of branches internal to the clade.

**Figure 2.**
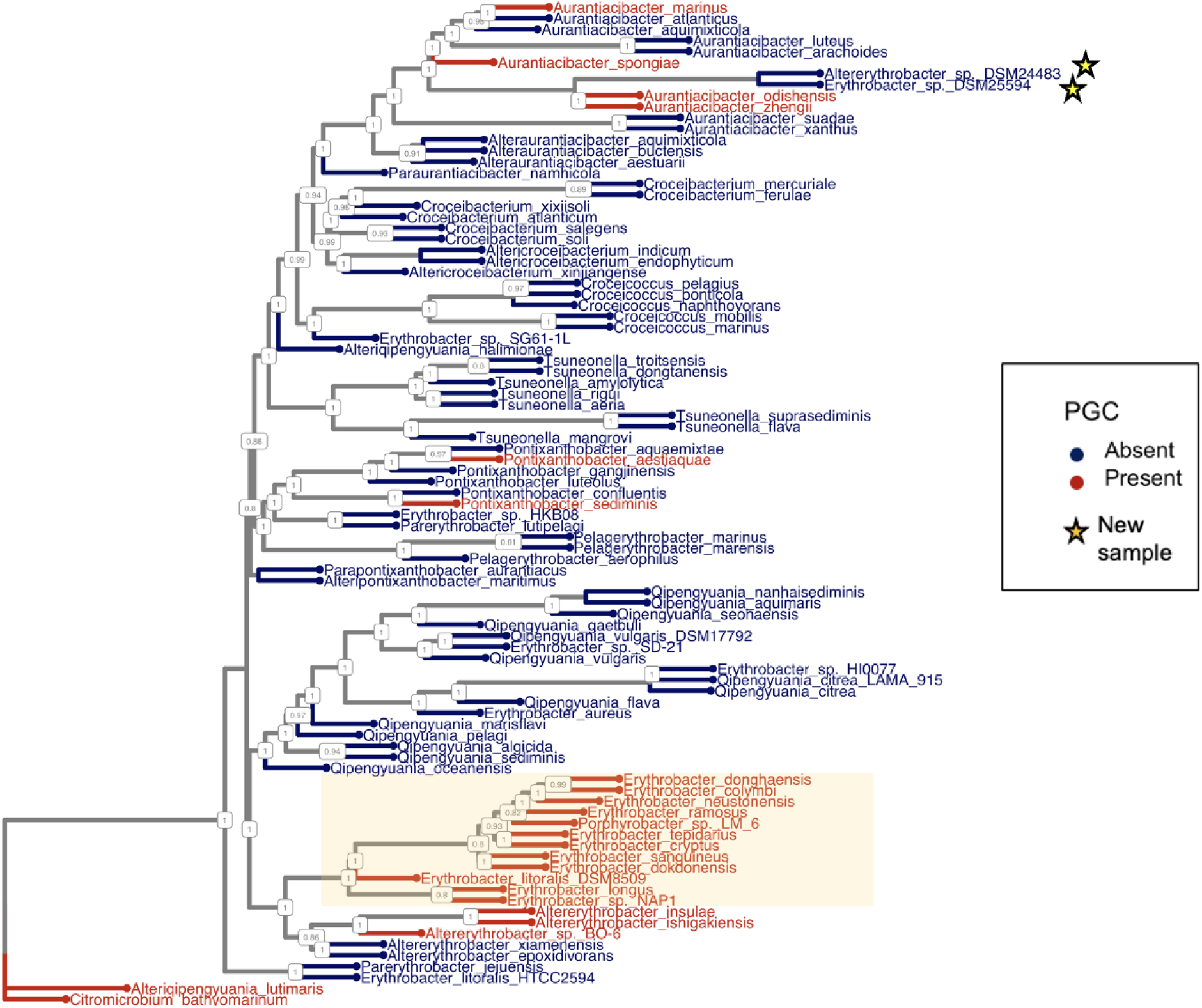
Phylogenomic tree of 90 *Erythrobacteraceae* isolates with *Citromicrobium bathyomarinum* as outgroup, inferred from 123 single-copy orthologs. Gene trees were inferred in RAxML with 100 bootstraps and a consensus species tree was constructed in ASTRAL-III. Support values are reported as local posterior probabilities. Highlighted box indicates monophyletic group which both possesses the photosynthetic gene cluster and expresses bacteriochlorophyll *a* in culture.

**Figure 3.**
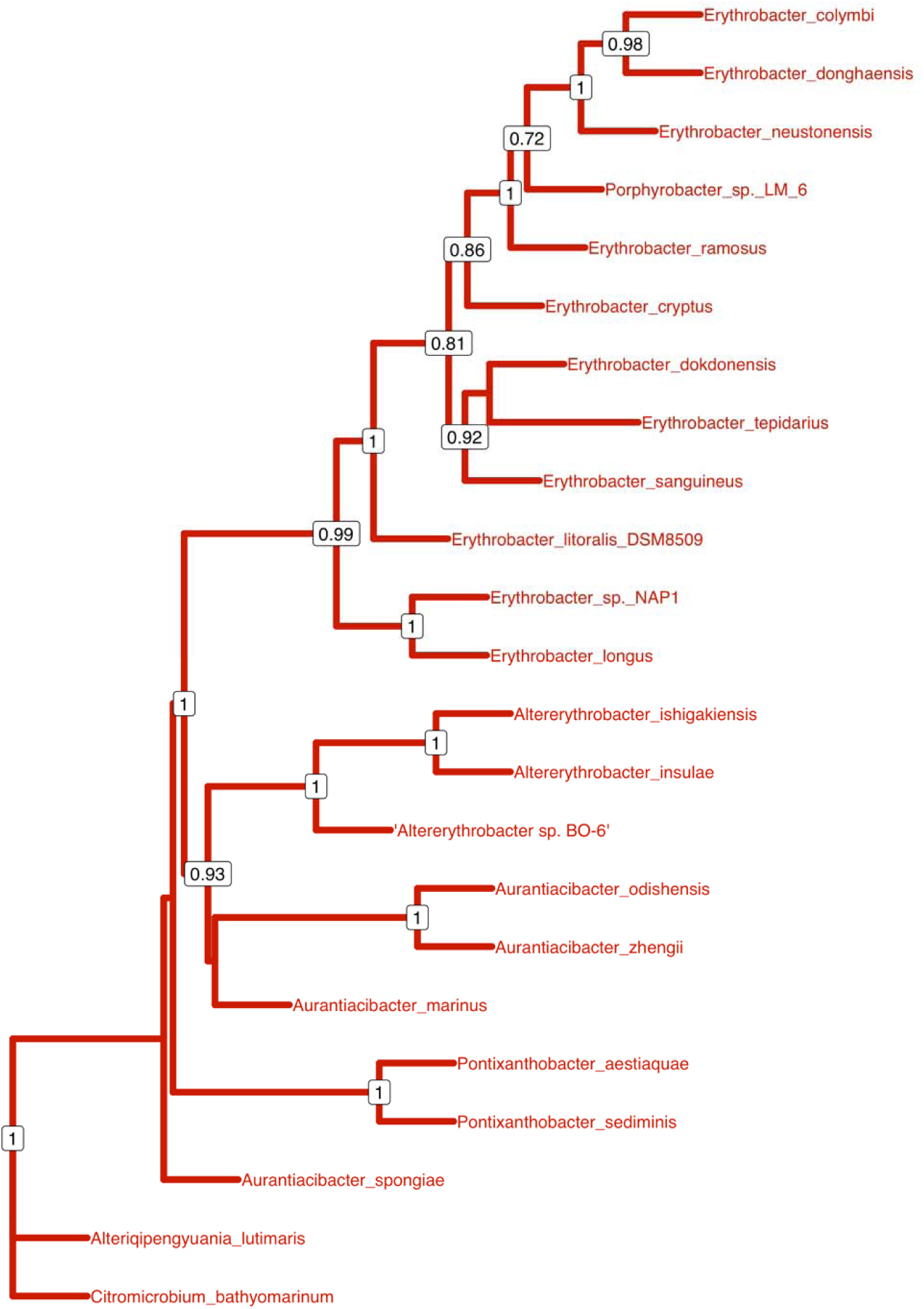
Phylogenetic tree for the photosynthetic gene cluster present in 23 *Erythrobacteraceae* isolates constructed from 25 genes conserved in >60% of taxa included in the dataset. Gene trees were inferred in RAxML with 100 bootstraps and a consensus species tree was constructed in ASTRAL-III. Support values are reported as local posterior probabilities.

### Ancestral State Reconstruction

Results of parsimony-based ancestral state reconstruction in Mesquite (**Figure 4)** indicate a most parsimonious scenario of at least four independent gains of the photosynthetic gene cluster across *Erythrobacteraceae*: for *Aurantiacibacter spongiae*, the branches for the sister taxa *Aurantiacibacter zhengii* and *Aurantiacibacter odishensis*, for *Pontixanthibacter aestiaquae*, and for *Pontixanthobacter sediminis*. There are two ambiguous steps at the root of the clade containing the genera *Erythrobacter* and *Altererythrobacter*, indicating two possible scenarios: either that the ancestor of the *Erythrobacter* and *Altererythrobacter* possessed the pohotosynthetic gene cluster, which was subsequently lost by the ancestors of *Altererythrobacter epoxidivorans* and *Altererythrobacter xiamenensis*, or alternatively, there were two independent gains, at the root of the *Erythrobacter* genus and the root of the *Altererythrobacter* sp. BO-6-*Altererythrobacter ishigakiensis-Altererythrobacter insulae* clade.

**Figure 4.**
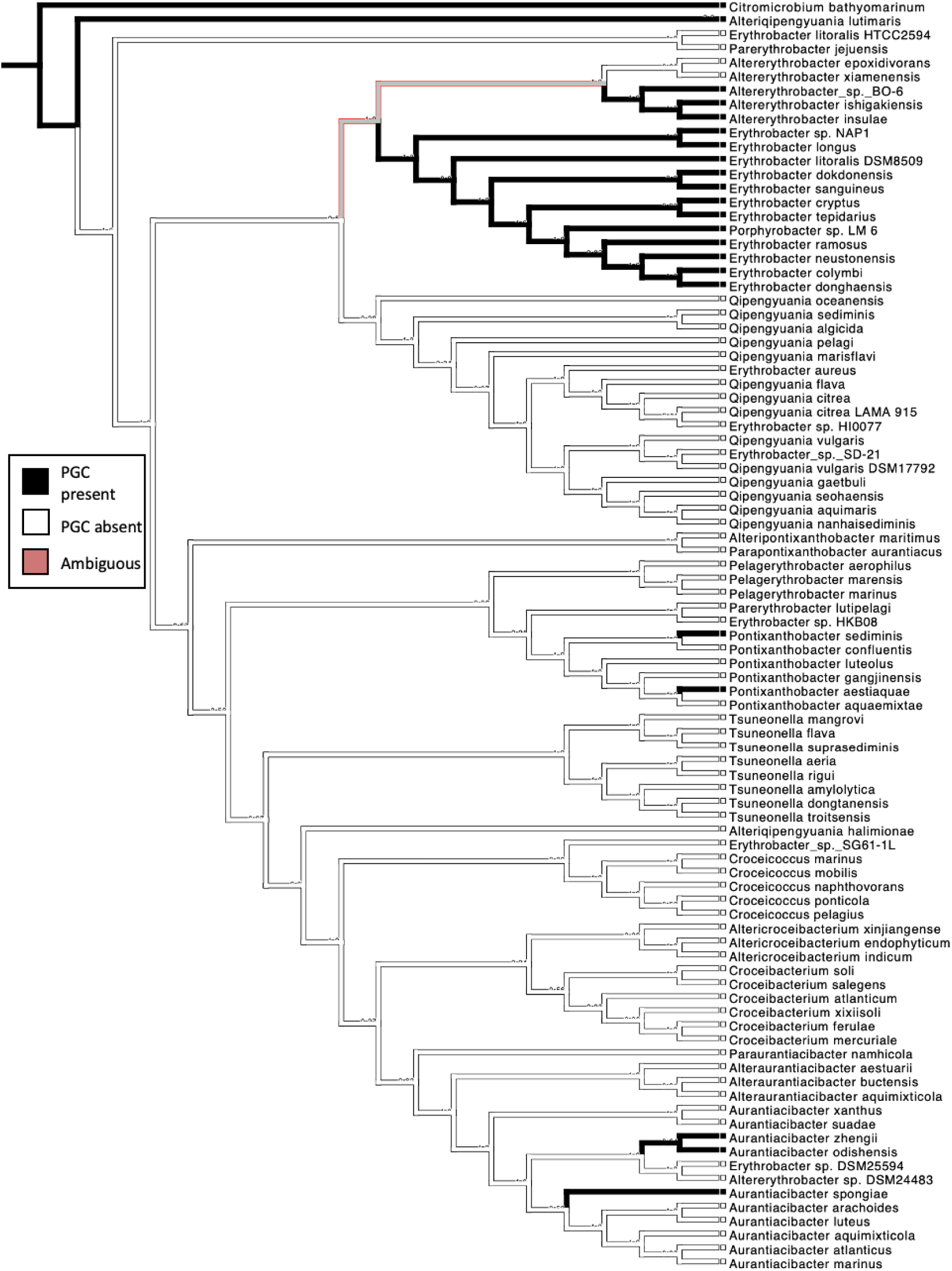
Parsimony-based ancestral state reconstruction for presence or absence of the photosynthetic gene cluster in *Erythrobacteraceae*. Presence of the PGC indicated in black, absence indicated in white, and ambiguous reconstruction indicated in red.

Maximum-likelihood-based reconstruction of ancestral states for *Erythrobacteraceae* (**Figure 5**) support independent gains of the photosynthetic gene cluster in *Aurantiacibacter spongiae*, *Pontixanthobacter aestiaquae,* and *Pontixanthobacter sediminis*. For the node at the root of *Aurantiacibacter zhengii, Aurantiacibacter odishensis, Erythrobacter* sp. DSM25594, and *Altererythrobacter* sp. DSM24483, the proportional likelihood of PGC presence is 0.786 and PGC absence is 0.214, at a ratio of 3.7:1, and is not significant based on the 7.4:1 ratio established by Schluter et al. (1997). At the root of *Erythrobacter* sp. DSM25594 and *Altererythrobacter* sp. DSM24483, the proportional likelihood of PGC absence is 0.986 and the likelihood of PGC presence is 0.014, a ratio of 70:1, providing significant evidence for PGC absence at this node. For the node at the root of the *Erythrobacter* and *Altererythrobacter* clades, the proportional likelihood of PGC absence is 0.866 and the likelihood of PGC presence is 0.134, a ratio of 6.5:1, below the threshold for significance; for the node at the root of the *Altererythrobacter* clade, the proportional likelihood of PGC absence is 0.872 and likelihood of PGC presence is 0.128, at a ratio of 6.8:1, also below the threshold for significance.

**Figure 5.**
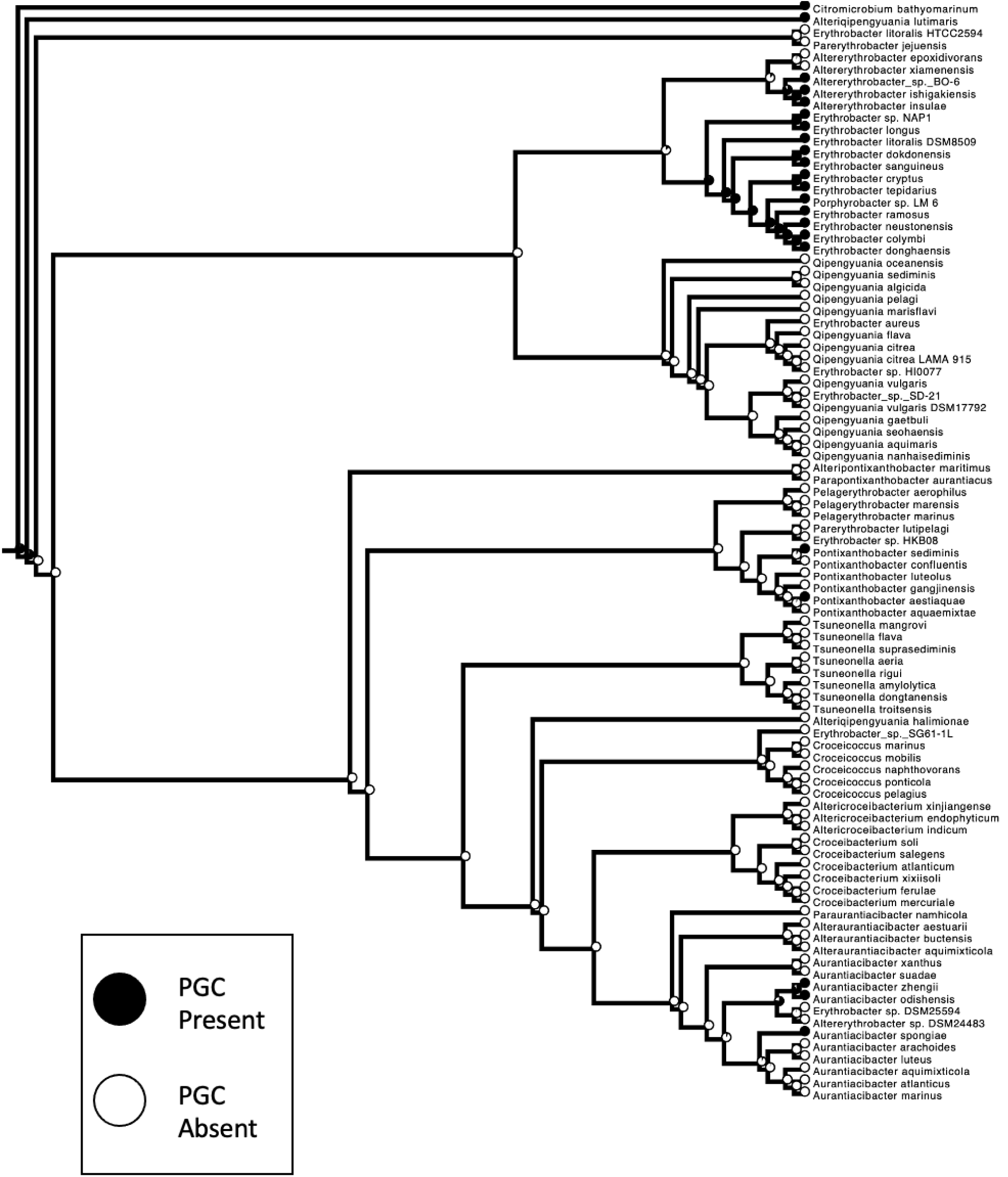
Maximum-likelihood ancestral state reconstruction for presence or absence of the photosynthetic gene cluster in *Erythrobacteraceae*. Presence of the PGC indicated in black, absence indicated in white.

### Multiple Genome Alignment of Photosynthetic Gene Cluster and Flanking Regions

Multiple genome alignment of photosynthetic-gene-cluster-bearing *Erythrobacteraceae* species revealed that the PGC is embedded in a diversity of integrative conjugative elements, ancestral plasmids which have integrated into the main chromosome of the ancestors of these species. In the large *Erythrobacter* clade whose members have documented patterns of expression of bacteriochlorophyll *a* in culture, the photosynthetic gene cluster is embedded in a conserved integrative conjugative element (ICE) across all twelve genomes (**Figure 6)**. This ICE was inserted just downstream of a tRNA-Arginine and contains a type IV secretion system conserved across all twelve taxa, encoding *TrbI/VirB10-TrbG/VirB9-VirB6-VirB4-VirB3-TrbC/VirB2*-lytic transglycosylase domain-containing protein-tetratricopeptide repeat protein. Downstream of the Type IV secretion system is a conserved transcription factor of the *OmpR/PhoB* family. Further downstream is a ferredoxin-NADP+ reductase, followed by three genes associated with iron transport: an Fe(3+) ABC transporter substrate-binding protein, an ABC transporter ATP-binding protein, and an iron ABC transporter permease. These final three genes are immediately followed by the *bchI* gene from the photosynthetic gene cluster across all twelve taxa.

**Figure 6.**
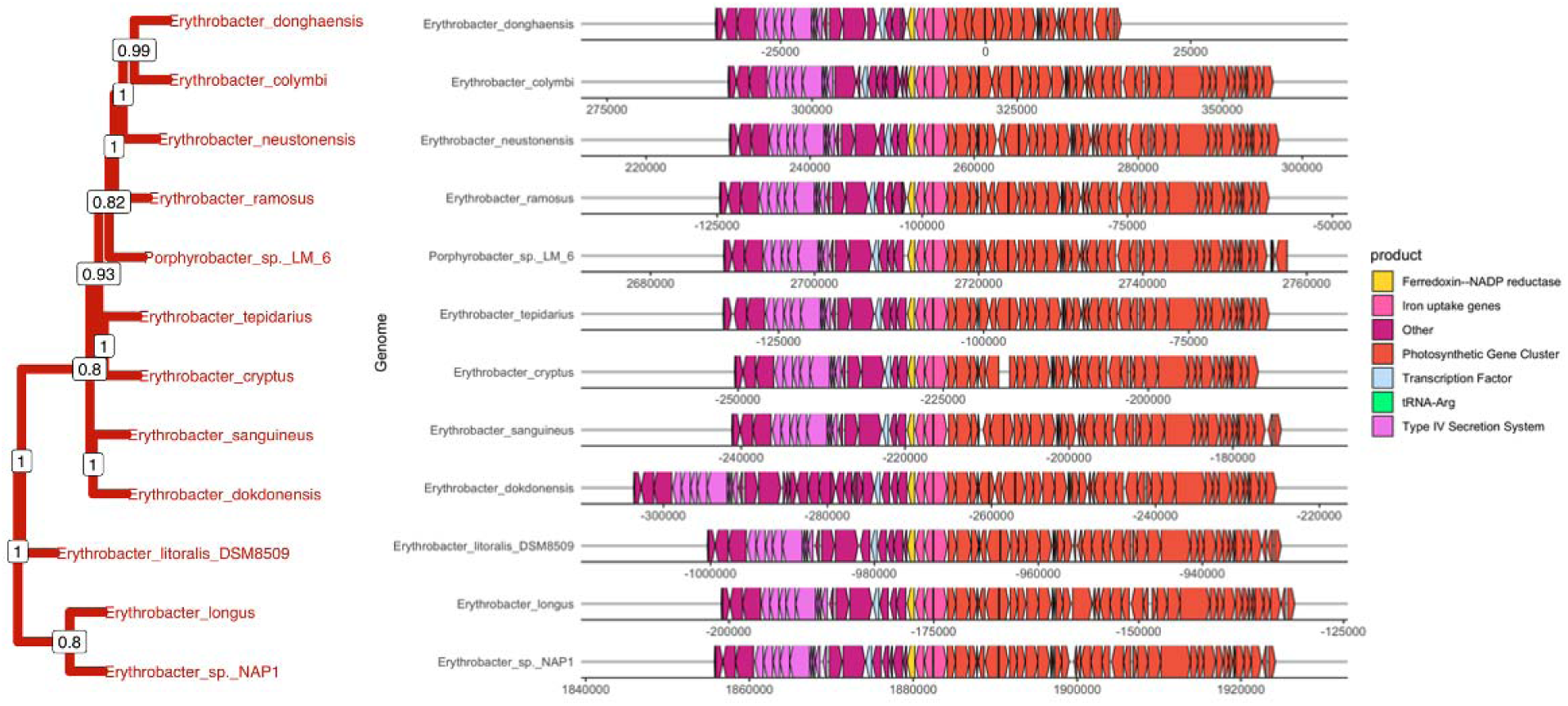
Multiple genome alignment for regions flanking the photosynthetic gene cluster in the *Erythrobacter* clade. Phylogeny trimmed from photosynthetic gene cluster tree.

In the *Altererythrobacter ishigakiensis-Altererythrobacter insulae-Altererythrobacter* sp. BO-6 clade, meanwhile, no such conserved integrative conjugative element is found (**Figure 7)**. The PGC of *Altererythrobacter insulae* and *Altererythrobacter ishigakiensis* is found immediately downstream of a conserved block of genes terminating in the gene for an SLC5 family protein similar to YidK in *Escherichia coli*, while in *Altererythrobacter sp.* BO-6 the PGC is downstream of a transcriptional regulator of the TetR/AcrR family. Downstream of the PGC in the genomes of both *Altererythrobacter ishigakiensis* and *Altererythrobacter* sp. BO-6 are two separate genomic islands not conserved between these two genomes.

**Figure 7.**
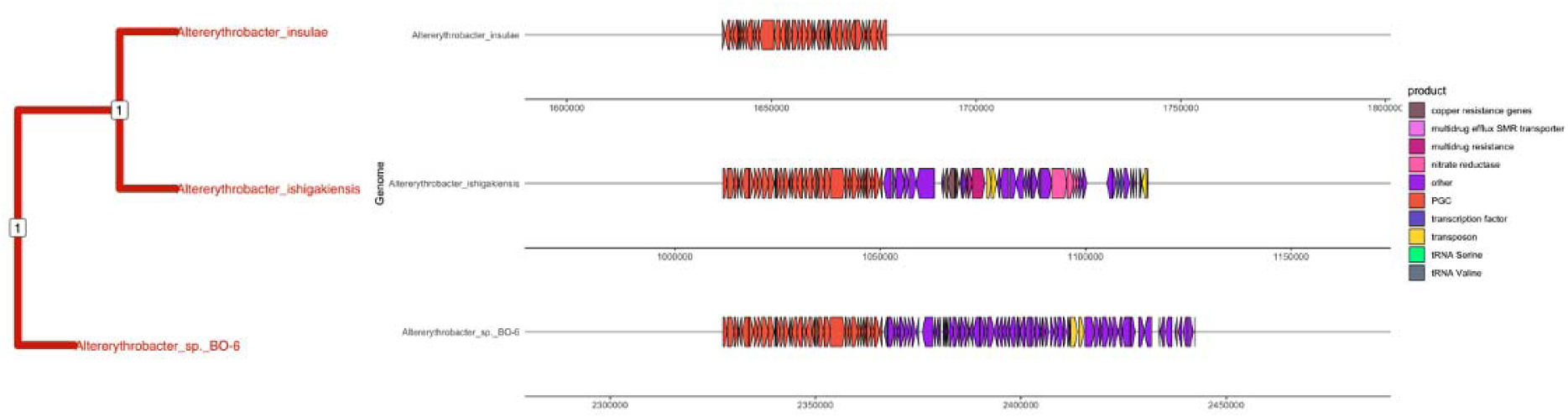
Multiple genome alignment for regions flanking the photosynthetic gene cluster in the *Altererythrobacter* sp. BO-6-*Altererythrobacter ishigakiensis-Altererythrobacter insulae* clade. Phylogeny trimmed from photosynthetic gene cluster tree as described in Chapter III.

Gene order data for the regions flanking the photosynthetic gene cluster reveal the superoperon is contained within diverse genomic islands among the remaining branches of the *Erythrobacteraceae* which possess it. In *Aurantiacibacter marinus*, the photosynthetic gene cluster is downstream of genes from a type IV secretion system and two genes annotated as coding for phage tail proteins (**Supplementary Figure 1**). In the genomes of sister species *Aurantiacibacter odishensis* and *Aurantiacibacter zhengii*, conserved elements of a genomic island upstream of the photosynthetic gene cluster in both species include genes for a cbb3-type cytochrome c oxidase, a gene annotated as oxygen-independent coproporphyrinogen III oxidase, and a cadmium-translocating P-type ATPase (**Supplementary Figure 2**). However, the genomic island containing *Aurantiacibacter zhengii* is much more extensive than in *Aurantiacibacter odishensis,* spanning 225,000 base pairs between the tRNA-Leucine and tRNA-Serine, and contains genes for a toxin-antitoxin system, for two complete type IV secretion systems, type I secretion system efflux RND transporter proteins, and multiple transposases and integrases. In *Aurantiacibacter spongiae*, the photosynthetic gene cluster’s internal gene order is inverted relative to the other PGCs present in *Erythrobacteraceae*, with a gene coding for 5-aminolevulinate and *puhE* following after *pufM*; the photosynthetic gene cluster is embedded in a genomic island which also contains genes for a cbb3-type cytochrome c oxidase (**Supplementary Figure 3**).

The photosynthetic gene cluster of *Pontixanthobacter sediminis* has been fragmented, with two segments separated by separated by 703,000 base pairs (**Supplementary Figure 4**): the first half of the photosynthetic gene cluster, spanning from a fragmentary copy of *bchI* to *pufM*, is on a genomic island downstream of tRNA-Lysine; no genes for conjugal transfer machinery are present adjacent to this half of the photosynthetic gene cluster. The second half of the photosynthetic gene cluster, spanning from a hypothetical protein conserved across *Erythrobacteraceae* PGCs to a gene for a cytochrome c family protein, is also present on a genomic island downstream of a tRNA-Leucine, which also contains genes for a type I restriction-modification system, a tyrosine-type recombinase/integrase, a transposase, and genes for a type II toxin-antitoxin system. In *Pontixanthobacter aestiaquae,* however, the photosynthetic gene cluster is downstream of a tRNA-Serine but is embedded within genes conserved among other members of *Erythrobacteraceae* (**Supplementary Figure 5**).

### Test for Relaxation of Selection

Results of the hypothesis test that selective pressure on the photosynthetic gene cluster has become relaxed on branches where the PGC is not expressed relative to those branches where bacteriochlorophyll *a* has been documented as expressed in culture, performed using the *RELAX* algorithm implemented in Hyphy, indicate no significant evidence for relaxation or intensification of selection among test branches relative to the reference branches (K = 0.97, p = 1.0000). The *RELAX* algorithm was also applied to individual genes from the photosynthetic gene cluster to test for heterogeneity in selective pressure across the PGC (**Table 1**). Five genes (*bchD, crtI, pufM, tspO, ppsR*) showed evidence for relaxation of selection at p = 0.05, while an additional five genes (*bchZ, pucC, bchN, puhB, acsF*) showed evidence for intensification of selection.

## DISCUSSION

While the aerobic anoxygenic photosynthesis was first described in *Erythrobacter longus* in 1979 by Shiba and Simudu, an account of how this mode of photosynthesis became patchily distributed throughout *Erythrobacteraceae* is still required (1982b). Findings in this paper for *Erythrobacteraceae* specifically are relevant to future investigations of the pattern of patchy distribution of AAP in other clades of Alphaproteobacteria, where similar techniques may further illuminate whether horizontal gene transfer is the primary explanation for the distribution of the photosynthetic gene cluster in *Erythrobacteraceae* specifically or in all clades where it is present.

The species phylogeny (**Figure 2**) for *Erythrobacteraceae,* inferred through phylogenomic methods in this study, consists of 17 identifiable clades, including the root, and largely agrees with previous phylogenetic work of Xu et al., with some key differences. Subclades reconstructed in the species tree correspond to the newly described genera of *Aurantiacibacter, Alteraurantiacibacter, Croceibacterium, Altericroceibacterium, Croceicoccus, Tsuneonella, Pontixanthobacter, Pelagerythrobacter, Qipengyuania, Erythrobacter,* and *Altererythrobacter*. The species tree inferred by Xu et al. was unrooted; for this study, the phylogeny was rooted with *Citromicrobium bathyomarinum*, and *Alteriqipengyuania lutimaris*, rather than clustering with *Alteriqipengyuamnia halimionae,* appeared most closely related to *C. bathyomarinum* at the root of the tree, thus challenging its family assignment as a member of *Erythrobacteraceae* altogether. Newly sequenced strains *Erythrobacter* sp. DSM25594 and *Altererythrobacter* sp. DSM24483 fall within the *Aurantiacibacter* clade. *Parapontixanthobacter aurantiacus* and *Alteripontixanthobacter maritimus* appear as sister taxa in this tree and should be assigned to a single genus. *Erythrobacter litoralis* HTCC2594, rather than being most closely related to *Erythrobacter litoralis* DSM8509 appears to be a sister species with *Parerythrobacter jejuensis* and falls entirely outside of the *Erythrobacter* clade. *Erythrobacter* sp. NAP1 and *Porphyrobacter* sp. LM 6 fall within the *Erythrobacter* clade, but several other *Erythrobacter* strains fall outside the *Erythrobacter* genus. *Erythrobacter* sp. HI0077 and *Erythrobacter* sp. SD-21 fall within the *Qipengyuania* clade. *Erythrobacter* sp. HKB08 is sister to *Parereythrobacter lutipelagi* – and *P. lutipelagi* is more closely related to the *Pontixanthobacter* clade than to *Parerythrobacter jejuensis.* Finally, *Erythrobacter* sp. SG61-1L is outgroup to the *Croceicoccus* clade. Thus, while the taxonomic revision of Xu et al. is largely supported by the phylogenomic work in this study, further taxonomic reassignment is necessary for the *Erythrobacteraceae* strains included in this phylogeny but not in previous work, and the *Parerythrobacter, Parapontixanthobacter, Alteripontixanthobacter,* and *Alteriqipengyuania* genera are not supported.

Consistent with the work of Xu et al., all species which both possess the photosynthetic gene cluster and express bacteriochlorophyll *a* in culture form one clade with a posterior probability of 1.0, the majority of which were assigned by Xu et al. to the *Erythrobacter* genus. This clade is recapitulated with a posterior probability of 0.99 in the tree inferred from 25 conserved genes from the photosynthetic gene cluster (**Figure 3**), although there are some rearrangements of branches within the clade between the two trees. The photosynthetic gene cluster phylogeny also groups *Alteriqipengyuania lutimaris* with *Citromicrobium bathyomarinum* at the root of the tree.

### *Ancestral state of the photosynthetic gene cluster in* Erythrobacteraceae

In this paper we have developed several lines of evidence that can be brought to bear on the question of whether the patchy distribution of the photosynthetic gene cluster in *Erythrobacteraceae* evolved due to a scenario of pure secondary loss of ancestral possession of the PGC, a scenario of pure gain through horizontal operon transfer, or a mixed regime of transfer and loss. Parsimony reconstructions support four independent gains for *Aurantiacibacter spongiae*, the sister species *Aurantiacibacter zhengii* and *Aurantiacibacter odishensis*, for *Pontixanthobacter aestiaquae*, and for *Pontixanthobacter* sediminis. Two steps are required to explain the pattern of PGC presence/absence in the *Erythrobacter-Altererythrobacter* clade: either two gains, at the root of *Erythrobacter* and at the root of *Altererythrobacter ishigakiensis, Altererythrobacter insulae*, and *Altererythrobacter* sp. BO-6, or else again at the root of *Erythrobacter* and *Altererythrobacter* and a loss by the ancestor of *Altererythrobacter epoxidivorans* and *Altereythrobacter xiamenensis*. In the maximum likelihood ancestral state reconstruction, the former scenario – of two gains – is more probable, but the relative likelihoods do not reach the level of significance. Additionally, the maximum likelihood reconstruction disagrees with the parsimony in one point by favoring a scenario of loss of the photosynthetic gene cluster from the ancestor of *Erythrobacter* sp. DSM25594 and *Altererythrobacter* sp. DSM24483, but the relative likelihoods again do not reach the level of significance.

Analysis of syntenic blocks flanking the photosynthetic gene cluster illuminates these ambiguities. For the first problem, whether two independent gains or a gain and a loss explain the pattern of PGC presence/absence in the *Erythrobacter-Altererythrobacter* clade, gene order data is clarifying. All genomes in the *Erythrobacter* clade possess a conserved integrative conjugative element of between 25,000 and 40,000 base pairs in length. This integrative conjugative element is absent from *Altererythrobacter ishigakiensis, Altererythrobacter insulae,* and *Altererythrobacter* sp. BO-6 (and indeed, from all other members of *Erythrobacteraceae* which possess the PGC), evidence for a gain of the photosynthetic gene cluster as part of a larger integrative conjugative element at the root of the *Erythrobacter* clade. This eliminates the scenario in which the photosynthetic gene cluster was gained at the root of *Erythrobacter* and *Altererythrobacter* and lost by *Altererythrobacter epoxidivorans* and *Altererythrobacter xiamenensis*. The photosynthetic gene cluster has been documented as acquired through an integrative conjugative element in the *Citromicrobium* genus, and integrative conjugative elements have been documented in *Erythrobacter* genomes previously, but integrative conjugative elements have not previously been documented as driving acquisition of the photosynthetic gene cluster in members of *Erythrobacteraceae* (Zheng et al., 2012, 2016). Examination of gene order data for regions flanking the photosynthetic gene cluster in other members of *Erythrobacteraceae* reveal no other conserved integrative conjugative elements or other flanking elements, except perhaps in the sister species *Aurantiacibacter odishensis* and *Aurantiacibacter zhengii*. The photosynthetic gene cluster is embedded on a large genomic island in *Aurantiacibacter zhengii*, which may be the result of repeated insertions between the tRNA-Leucine and tRNA-Serine, and which is mostly absent from *Aurantiacibacter odishensis*, except for genes for a cbb3-type cytochrome c oxidase, a gene annotated as oxygen-independent coproporphyrinogen III oxidase, and a cadmium-translocating P-type ATPase, which are conserved not only in content but also in gene order and position adjacent to the photosynthetic gene cluster. We hypothesize that the photosynthetic gene cluster was acquired through horizontal operon transfer by the ancestor of *Aurantiacibacter zhengii* and *Aurantiacibacter odishensis*, and that *Aurantiacibacter zhengii* subsequently experienced the integration of additional genetic material through horizontal transfer at the same genomic hotspot.

The evidence thus tends toward a conclusion that horizontal operon transfer, rather than secondary loss or a mixed regime of loss and transfer, drove the acquisition and distribution of the photosynthetic gene cluster in *Erythrobacteraceae*.

### Selective pressure on the photosynthetic gene cluster across the phylogeny

While results of the RELAX test on concatenated gene sequences from the photosynthetic gene cluster do not support the hypothesis that selective pressure has become relaxed in PGC-bearing members of *Erythrobacteraceae* outside the *Erythrobacter* clade, results of the RELAX test on individual genes from the PGC reveal a more heterogeneous picture. Five genes (*bchD, crtI, pufM, tspO, ppsR*) showed evidence for relaxation of selection, while an additional five genes (*bchZ, pucC, bchN, puhB, acsF*) showed evidence for intensification of selection.

Genes which play a role in the biosynthesis of bacteriochlorophyll *a* show evidence both for relaxation of selection (*bchD, tspO*) and intensification of selection (*bchZ, bchN, acsF);* a similar pattern is seen for genes whose protein products either form components of the photosynthetic reaction center (*pufM*, evidence for relaxation of selection) or are involved in reaction center assembly (*puhB,* evidence for intensification of assembly). These contradictory signals, combined with the lack of evidence for relaxation of selection in a concatenated alignment of genes from the photosynthetic gene cluster, lead us to reject the hypothesis that selection is relaxed in the photosynthetic gene cluster in non-*Erythrobacter* branches relative to the *Erythrobacter* clade. However, that the photosynthetic gene cluster is expressed in culture in only some clades where the PGC is present does not mean that it is not expressed in the environment in the other clades; this may explain why selection would be maintained on genes that are not expressed in culture.

### Conclusions

In this paper we generate evidence supporting multiple gains of the PGC in *Erythrobacteraceae* by horizontal gene transfer mediated by integrative conjugative elements, consistent with prior work on the photosynthetic gene cluster in *Rhodobacteraceae*, where plasmid-mediated transfer of the PGC was documented (Brinkmann et al., 2018). While no evidence of secondary loss of the PGC was found, we cannot exclude this possibility given the incompleteness of any phylogeny for bacteria relative to all possible existing strains. Notably, only one clade within *Erythrobacteraceae* has been documented to both possess the photosynthetic gene cluster within its genome and to express that gene cluster, the clade corresponding to the *Erythrobacter* genus. The photosynthetic gene cluster on that clade is embedded in a conserved integrative conjugative element which also contains several other genes not previously documented as part of the photosynthetic gene cluster, raising the question of whether those genes may enable the expression of the PGC in *Erythrobacter*. Regardless, aerobic anoxygenic photosynthesis in *Erythrobacteraceae* presents researchers with an unanswered question: why is the photosynthetic gene cluster not consistently expressed across the family? Possible explanations for future investigation include that the PGC is expressed *in natura* but not in culture in other clades of *Erythrobacteraceae*, or that other regulatory elements or genes are necessary for expression of photosynthesis beyond simply the conserved suite of genes on the PGC.

