## Supplementary Figures and Tables for "The Patchy Distribution Of The Photosynthetic Gene Cluster In *Erythrobacteraceae*"

**Supplementary Figure 1**. Plasmid of 61,937 base pairs present in genomes of both strains DSM24483 and DSM25594. Pink: transposase genes; Orange: type II toxin-antitoxin system genes; Dark green: ParA/ParB chromosome partitioning protein genes; Blue: cytochrome c oxidase genes; Teal: molybdenum cofactor biosynthesis genes; Maroon: nitrite transport and reduction genes; Light green: type IV secretion system genes.

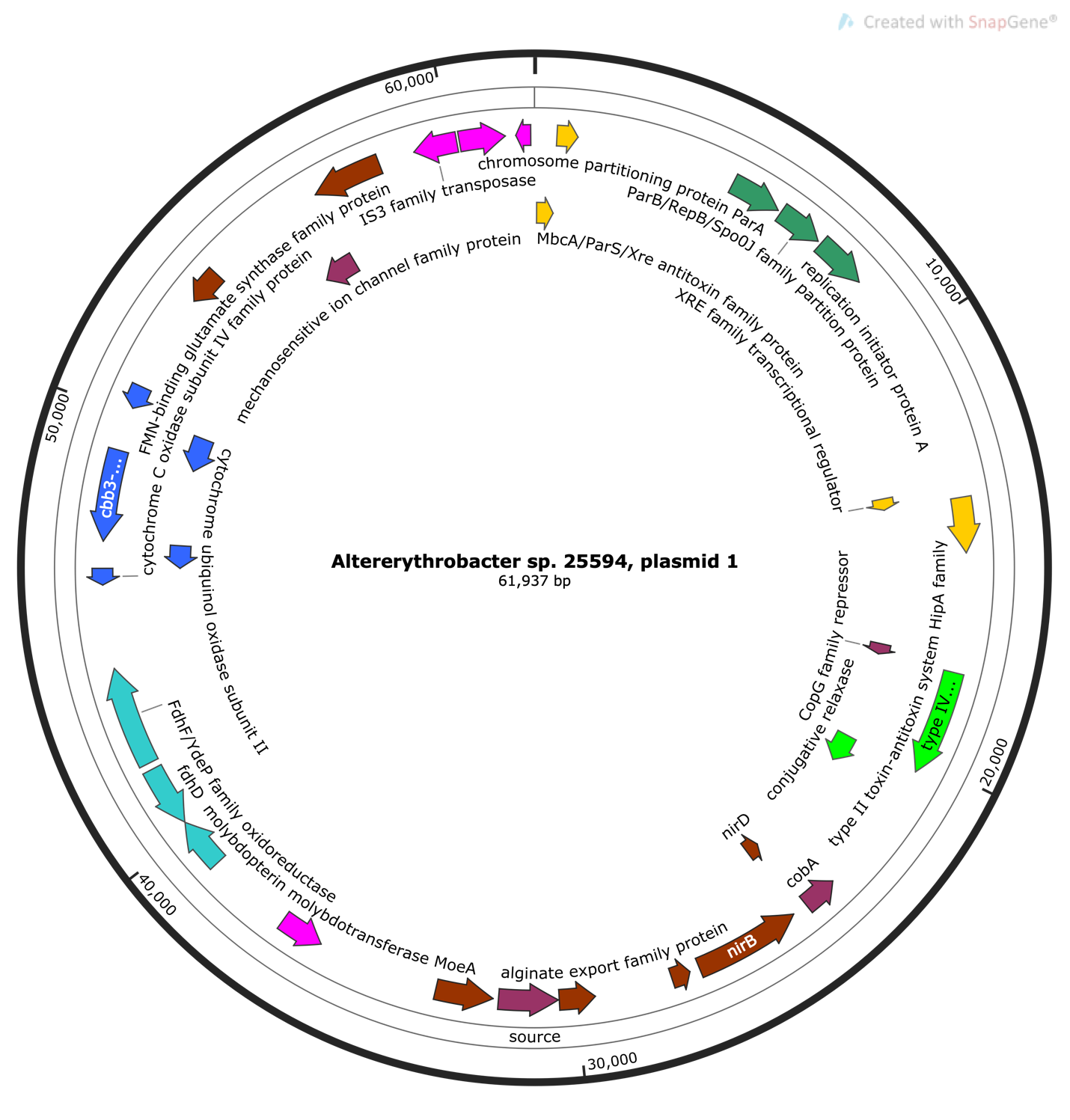

**Supplementary Figure 2.** Plasmid of 35,415 base pairs present in both strains DSM24483 and DSM25594. Pink: transposase genes; Dark green: ParA/ParB genes; Light green: type IV secretion system; Purple: metalloregulator ArsR/SmtB repressor and cation diffusion facilitator family transporter.

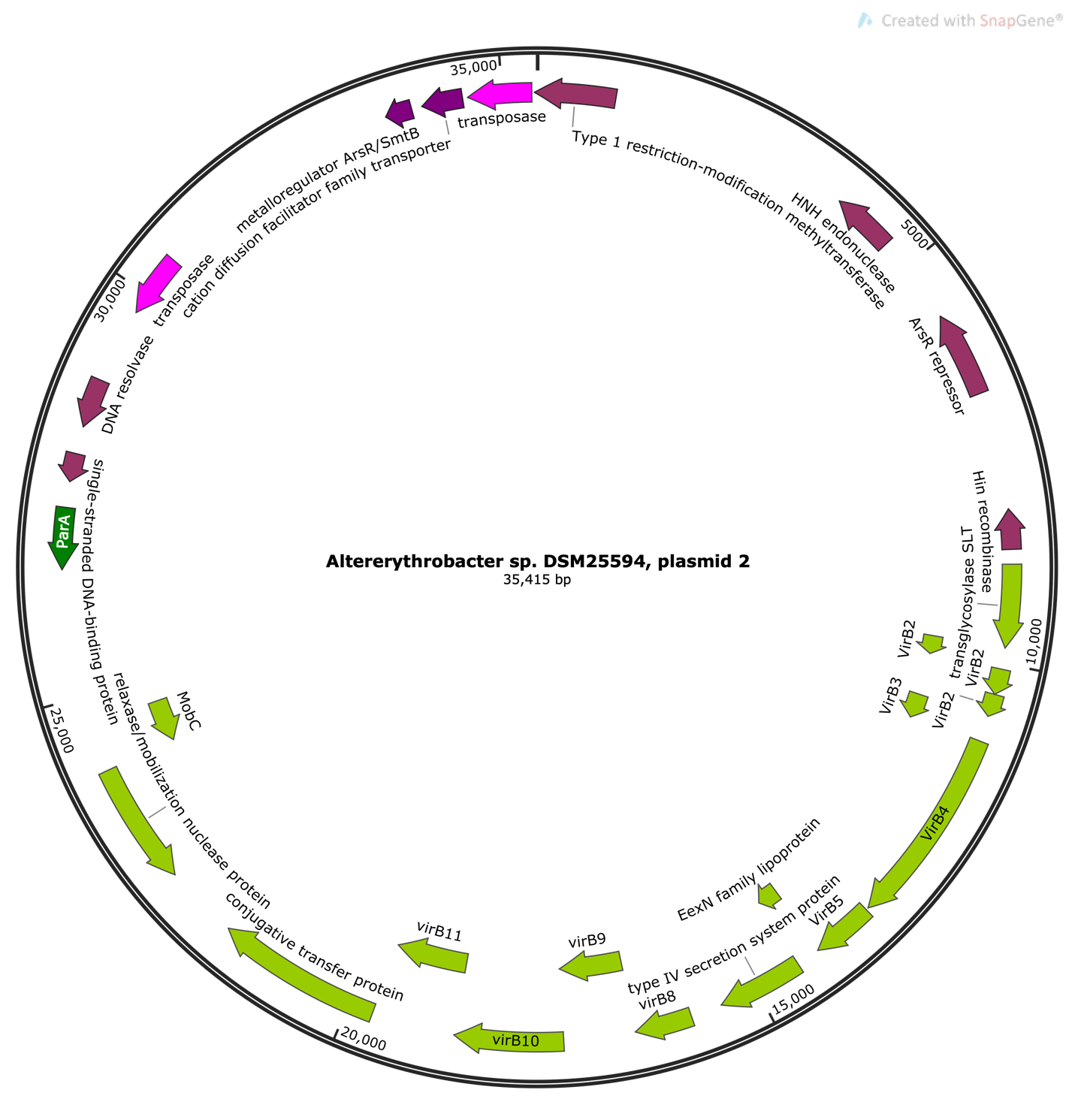

**Supplementary Figure 3.** Plasmid of 24,880 base pairs present in both strains DSM24483 and DSM25594. Dark green: Replication and partitioning genes. Red: endonuclease genes. Aqua: Copper translocation and reduction genes. Blue: cytochrome C oxidase gene. Purple: cation diffusion facilitator family transporters. Light green: type IV secretion system genes.

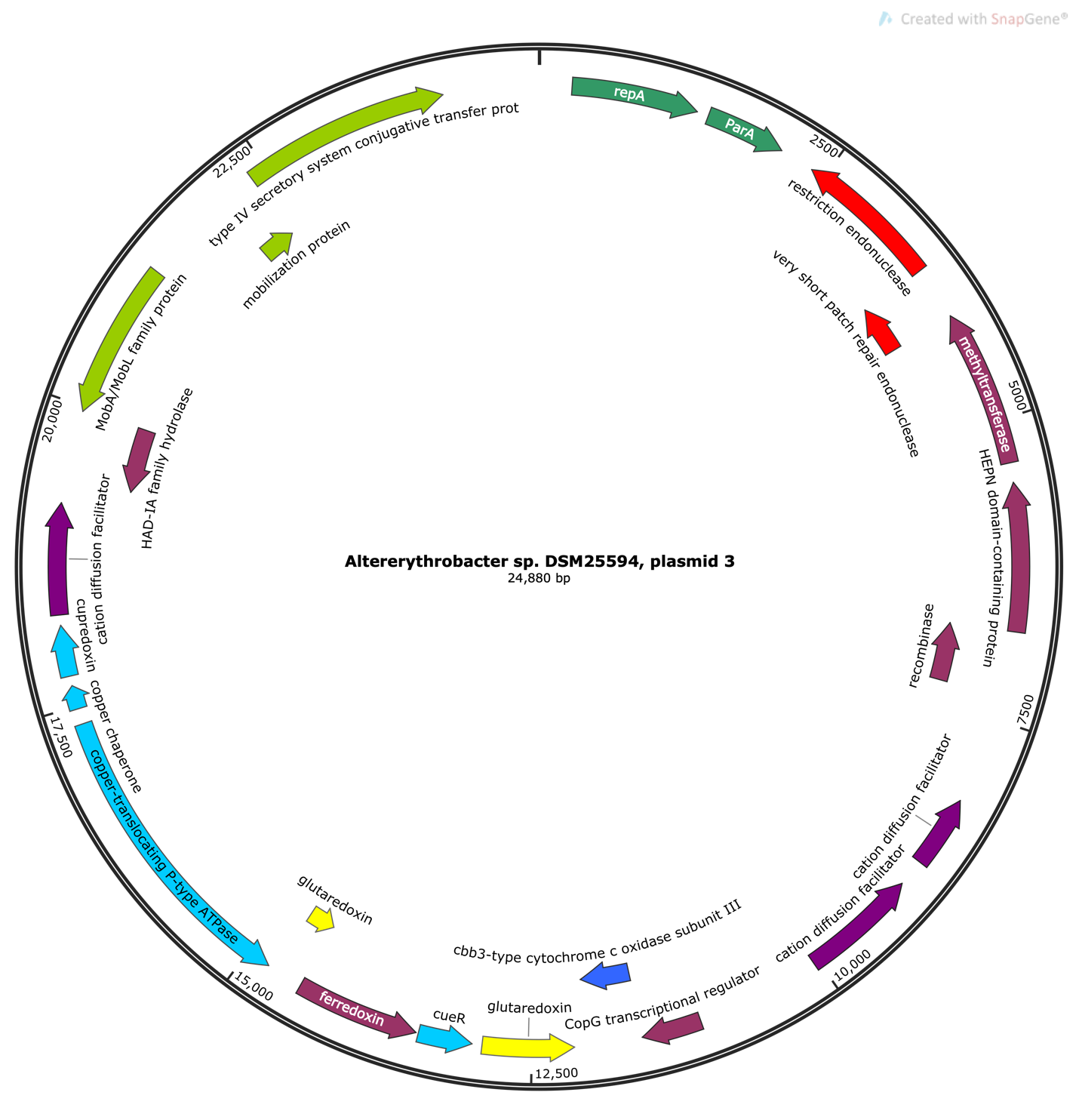

**Supplementary Figure 4.** Genomic island of 47,514 base pairs from *Altererythrobacter* sp. DSM24483. Light green: Type IV secretion system conjugal transfer machinery. Blue: cytochrome p450 genes. Pink: transposases.

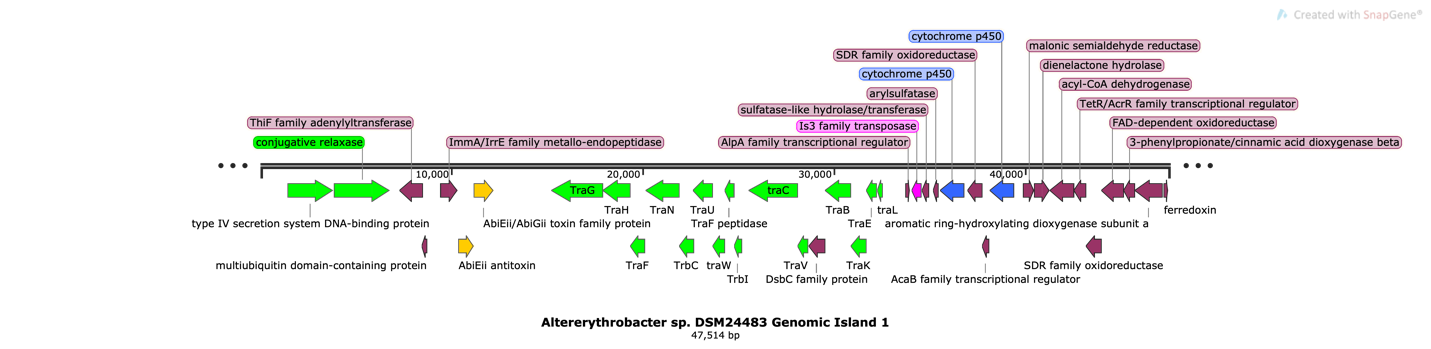

**Supplementary Figure 5.** Genomic island of 20,952 base pairs from *Altererythrobacter* sp. DSM24483. Green: tRNA for Threonine.

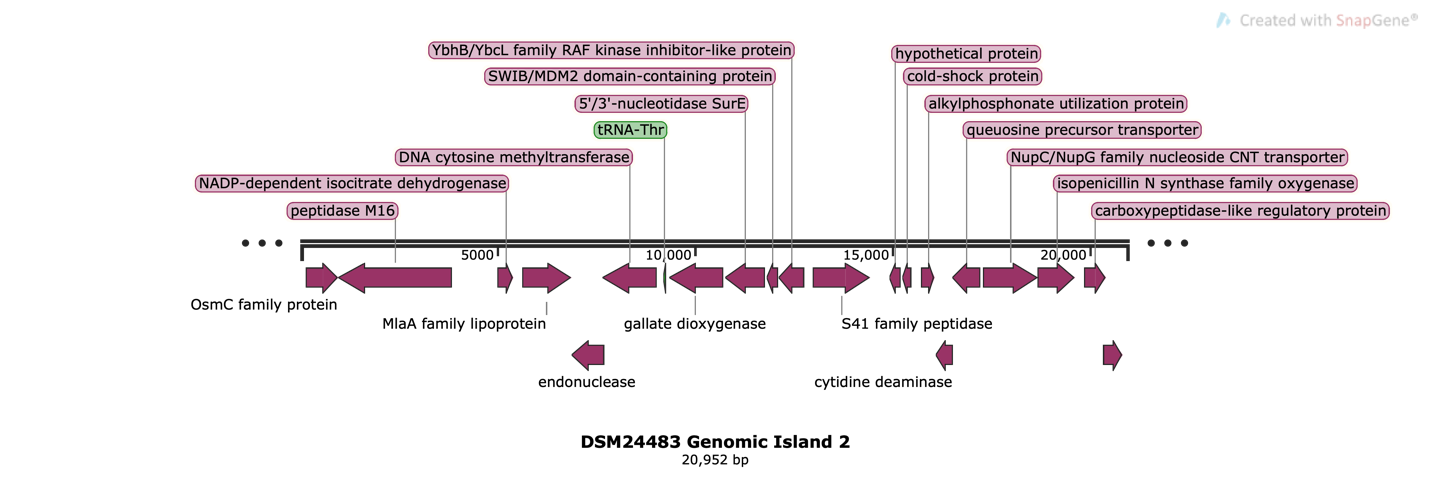

**Supplementary Figure 6.** Genomic Island of 44,296 base pairs from *Altererythrobacter* sp. DSM24483. Yellow: Type II secretion system genes. Light pink: Type I secretion system genes. Slate blue: *ScpA/ScpB* segregation/condensation proteins. Salmon pink: *TatA/TatB/TatC* translocation protein transport proteins.

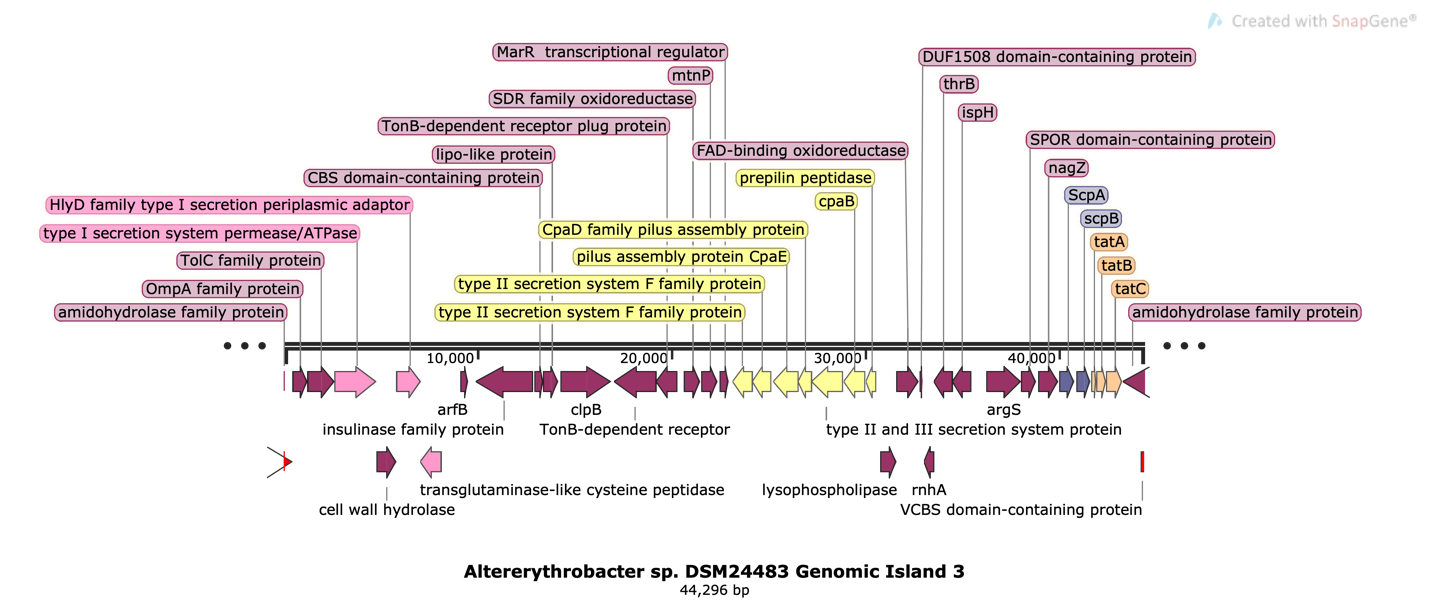

**Supplementary Figure 7.** Genomic island of 3,770 base pairs in *Altererythrobacter* sp. DSM24483. Pink: transposase.

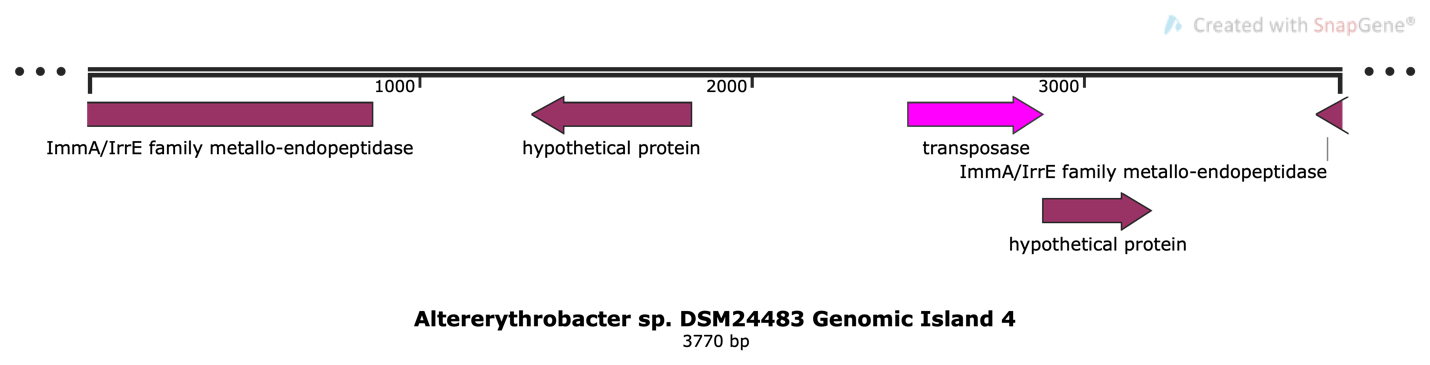

**Supplementary Table 1.** Accessions for genomes included in phylogenomic analysis, along with current and previous specific names, whether they were included in previous phylogenomic work by Xu *et al* 2020, and whether they possess and express the photosynthetic gene cluster in culture.

| Accession | Species Name | Previous Name | Included in Xu et al? | Possesses Photosynthetic Gene Cluster | Expresses Bacteriochlorophyll *a* in culture |
| --- | --- | --- | --- | --- | --- |
| N/A | *Erythrobacter* sp. DSM25594 | N/A | No | No | No |
| N/A | *Altererythrobacter* sp. DSM24483 | N/A | No | No | No |
| NC_007722 | *Erythrobacter litoralis* HTCC2594 | N/A | No | No | No |
| NZ_AAMW01000004 | *Erythrobacter* sp. NAP1 | N/A | No | Yes | Yes |
| NZ_ABCG01000001 | *Erythrobacter* sp. SD-21 | N/A | No | No | No |
| NZ_ADAE01000001 | *Citromicrobium bathyomarinum* | N/A | No | Yes | Yes |
| NZ_AUHC01000001 | *Erythrobacter cryptus* | *Porphyrobacter cryptus* | Yes | Yes | Yes |
| NZ_CCSI01000001 | *Qipengyuania vulgaris* | *Erythrobacter vulgaris* | No | No | No |
| NZ_CP011310 | *Aurantiacibacter atlanticus* | *Erythrobacter atlanticus* | Yes | No | No |
| NZ_CP011452 | *Croceibacterium atlanticum* | *Altererythrobacter atlanticus* | Yes | No | No |
| NZ_CP011770 | *Croceicoccus naphthovorans* | N/A | Yes | No | No |
| NZ_CP011805 | *Pelagerythrobacter marensis* | N/A | Yes | No | No |
| NZ_CP012669 | *Altererythrobacter epoxidivorans* | N/A | Yes | No | No |
| NZ_CP015963 | *Altererythrobacter ishigakiensis* | N/A | Yes | Yes | No |
| NZ_CP016033 | *Erythrobacter neustonensis* | *Porphyrobacter neustonensis* | Yes | Yes | Yes |
| NZ_CP016545 | *Paraurantiacibacter namhicola* | *Altererythrobacter namhicola* | Yes | No | No |
| NZ_CP016591 | *Tsuneonella dongtanensis* | *Altererythrobacter dongtanensis* | Yes | No | No |
| NZ_CP017057 | *Erythrobacter litoralis* DSM8509 | N/A | Yes | Yes | Yes |
| NZ_CP017113 | *Porphyrobacter* sp. LM 6 | N/A | No | Yes | Yes |
| NZ_CP019602 | *Croceicoccus marinus* | N/A | Yes | No | No |
| NZ_CP022528 | *Qipengyuania flava* | *Erythrobacter flavus* | Yes | No | No |
| NZ_CP022889 | *Tsuneonella mangrovi* | *Altererythrobacter mangrovi* | Yes | No | No |
| NZ_CP024920 | *Qipengyuania seohaensis* | *Erythrobacter seohaensis* | Yes | No | No |
| NZ_CP031357 | *Erythrobacter aureus* | N/A | No | No | No |
| NZ_CP032570 | *Tsuneonella amylolytica* | N/A | Yes | No | No |
| NZ_CP035310 | *Erythrobacter* sp. HKB08 | N/A | No | No | No |
| NZ_CP037948 | *Qipengyuania sediminis* | N/A | Yes | No | No |
| NZ_CP049259 | *Altererythrobacter* sp. BO-6 | N/A | No | Yes | No |
| NZ_FOWZ01000001 | *Qipengyuania nanhaisediminis* | *Erythrobacter nanhaisediminis* | Yes | No | No |
| NZ_FRDF01000001 | *Erythrobacter sanguineus* | *Porphyrobacter sanguineus* | Yes | Yes | Yes |
| NZ_FXWG01000001 | *Altererythrobacter xiamenensis* | N/A | Yes | No | No |
| NZ_JMIW01000001 | *Erythrobacter longus* | N/A | Yes | Yes | Yes |
| NZ_JTDN01000001 | *Croceibacterium mercuriale* | N/A | Yes | No | No |
| NZ_JXQC01000001 | *Erythrobacter* sp. SG61-1L | N/A | No | No | No |
| NZ_JYNE01000001 | *Qipengyuania citrea* | *Erythrobacter citreus* | Yes | No | No |
| NZ_LBHB01000001 | *Aurantiacibacter luteus* | *Altererythrobacter luteus* | Yes | No | No |
| NZ_LDCP01000001 | *Aurantiacibacter marinus* | *Altererythrobacter marinus* | Yes | Yes | No |
| NZ_LMAU01000001 | *Tsuneonella troitsensis* | *Altererythrobacter troitensis* | Yes | No | No |
| NZ_LSJJ01000001 | *Erythrobacter donghaensis* | *Porphyrobacter donghaensis* | Yes | Yes | Yes |
| NZ_LWFR01000001 | *Erythrobacter* sp. HI0077 | N/A | No | No | No |
| NZ_LYWY01000001 | *Croceicoccus pelagius* | N/A | Yes | No | No |
| NZ_LYWZ01000001 | *Croceicoccus mobilis* | N/A | Yes | No | No |
| NZ_LZYB01000001 | *Erythrobacter dokdonensis* | *Porphyrobacter dokdonensis* | Yes | Yes | Yes |
| NZ_MUYJ01000001 | *Erythrobacter tepidarius* | *Porphyrobacter tepidarius* | Yes | Yes | Yes |
| NZ_MUYK01000001 | *Erythrobacter colymbi* | *Porphyrobacter colymbi* | Yes | Yes | Yes |
| NZ_PHSO01000001 | *Tsuneonella flava* | *Altererythrobacter flavus* | Yes | No | No |
| NZ_QBKA01000001 | *Alteripontixanthobacter maritimus* | *Altererythrobacter maritimus* | Yes | No | No |
| NZ_QRBB01000010 | *Alteriqipengyuania lutimaris* | *Altererythrobacter lutimaris* | Yes | Yes | No |
| NZ_QURJ01000001 | *Altererythrobacter insulae* | *Altererythrobacter insulae* | Yes | Yes | No |
| NZ_QXFK01000014 | *Pelagerythrobacter aerophilus* | N/A | Yes | No | No |
| NZ_QXFL01000001 | *Aurantiacibacter zhengii* | *Erythrobacter zhengii* | Yes | Yes | No |
| NZ_QXFM01000001 | *Aurantiacibacter xanthus* | *Erythrobacter xanthus* | Yes | No | No |
| NZ_QYOS01000001 | *Aurantiacibacter odishensis* | *Erythrobacter odishensis* | Yes | Yes | No |
| NZ_QZVQ01000001 | *Croceibacterium ferulae* | N/A | Yes | No | No |
| NZ_RAHJ01000001 | *Tsuneonella suprasediminis* | *Altererythrobacter* sp. Ery12 | No | No | No |
| NZ_RAHX01000001 | *Aurantiacibacter aquimixticola* | *Erythrobacter aquimixticola* | Yes | No | No |
| NZ_RPFZ01000001 | *Aurantiacibacter spongiae* | N/A | Yes | Yes | No |
| NZ_RSEK01000001 | *Altericroceibacterium xinjiangense* | *Altererythrobacter xinjiangensis* | Yes | No | No |
| NZ_RSEL01000001 | *Tsuneonella rigui* | *Altererythrobacter rigui* | Yes | No | No |
| NZ_RXOL01000001 | *Croceicoccus ponticola* | N/A | No | No | No |
| NZ_SKCG01000001 | *Aurantiacibacter suaedae* | *Erythrobacter suaedae* | Yes | No | No |
| NZ_SKCJ01000001 | *Parerythrobacter lutipelagi* | *Altererythrobacter lutipelagi* | No | No | No |
| NZ_SSHH01000001 | *Alteraurantiacibacter aquimixticola* | *Altererythrobacter aquimixticola* | No | No | No |
| NZ_VCAO01000001 | *Qipengyuania marisflavi* | N/A | Yes | No | No |
| NZ_WTYA01000001 | *Qipengyuania algicida* | *Porphyrobacter algicida* | Yes | No | No |
| NZ_WTYB01000001 | *Erythrobacter ramosus* | *Erythromicrobium ramosum* | Yes | Yes | Yes |
| NZ_WTYC01000001 | *Qipengyuania vulgaris* DSM17792 | *Erythrobacter vulgaris* DSM17792 | Yes | No | No |
| NZ_WTYD01000001 | *Qipengyuania pelagi* | *Erythrobacter pelagi* | Yes | No | No |
| NZ_WTYE01000001 | *Parerythrobacter jejuensis* | *Erythrobacter jejuensis* | Yes | No | No |
| NZ_WTYF01000001 | *Qipengyuania gaetbuli* | *Erythrobacter gaetbuli* | Yes | No | No |
| NZ_WTYG01000010 | *Qipengyuania citrea* | N/A | Yes | No | No |
| NZ_WTYH01000001 | *Aurantiacibacter arachoides* | *Erythrobacter arachoides* | Yes | No | No |
| NZ_WTYI01000001 | *Qipengyuania aquimaris* | *Erythrobacter aquimaris* | Yes | No | No |
| NZ_WTYJ01000001 | *Croceibacterium xixiisoli* | *Altererythrobacter xixiisoli* | Yes | No | No |
| NZ_WTYK01000001 | *Croceibacterium soli* | *Altererythrobacter soli* | Yes | No | No |
| NZ_WTYL01000001 | *Pontixanthobacter sediminis* | *Altererythrobacter sediminis* | Yes | Yes | No |
| NZ_WTYM01000001 | *Croceibacterium salegens* | *Altererythrobacter salegens* | Yes | No | No |
| NZ_WTYN01000001 | *Qipengyuania oceanensis* | *Altererythrobacter oceanensis* | Yes | No | No |
| NZ_WTYO01000001 | *Pelagerythrobacter marinus* | *Altererythrobacter marinus* | Yes | No | No |
| NZ_WTYP01000001 | *Pontixanthobacter luteolus* | *Altererythrobacter luteolus* | Yes | No | No |
| NZ_WTYQ01000001 | *Altericroceibacterium indicum* | *Altererythrobacter indicus* | Yes | No | No |
| NZ_WTYR01000001 | *Alteriqipengyuania halimionae* | *Altererythrobacter halimionae* | Yes | No | No |
| NZ_WTYS01000001 | *Pontixanthobacter gangjinensis* | *Altererythrobacter gangjinensis* | Yes | No | No |
| NZ_WTYT01000001 | *Altericroceibacterium endophyticum* | *Altererythrobacter endophyticus* | Yes | No | No |
| NZ_WTYU01000001 | *Pontixanthobacter confluentis* | *Altererythrobacter confluentis* | Yes | No | No |
| NZ_WTYV01000001 | *Alteraurantiacibacter buctensis* | *Altererythrobacter buctensis* | Yes | No | No |
| NZ_WTYW01000001 | *Parapontixanthobacter aurantiacus* | *Altererythrobacter aurantiacus* | Yes | No | No |
| NZ_WTYX01000001 | *Pontixanthobacter aquaemixtae* | *Altererythrobacter aquaemixtae* | Yes | No | No |
| NZ_WTYY01000001 | *Alteraurantiacibacter aestuarii* | *Altererythrobacter aestuarii* | Yes | No | No |
| NZ_WTYZ01000001 | *Pontixanthobacter aestiaquae* | *Altererythrobacter aestiaquae* | Yes | Yes | No |
| NZ_WTZA01000001 | *Tsuneonella aeria* | *Altererythrobacter aerius* | Yes | No | No |

**Supplementary Table 2.** Genes included in phylogeny for the photosynthetic gene cluster across 23 taxa.

| **Gene Annotation** | **Present in % of Taxa** |
| --- | --- |
| bchI magnesium chelatase ATPase subunit I | 95.65% |
| magnesium chelatase subunit D | 95.65% |
| hydroxyneurosporene dehydrogenase | 100.00% |
| chlorophyll synthesis pathway protein BchC | 100.00% |
| chlorophyllide a reductase iron protein subunit X | 100.00% |
| chlorophyllide a reductase subunit Y | 100.00% |
| chlorophyllide a reductase subunit Z | 100.00% |
| light harvesting protein | 100.00% |
| light harvesting protein | 100.00% |
| photosynthetic reaction center subunit L | 100.00% |
| photosynthetic reaction center subunit M | 100.00% |
| geranylgeranyl diphosphate reductase | 100.00% |
| BCD family MFS transporter | 100.00% |
| chlorophyll synthase ChlG | 100.00% |
| 2-vinyl bacteriochlorophyllide hydratase | 100.00% |
| ferredoxin:protochlorophyllide reductase (ATP-dependent) subunit N | 100.00% |
| ferredoxin:protochlorophyllide reductase (ATP-dependent) subunit B | 100.00% |
| magnesium chelatase subunit H | 91.30% |
| ferredoxin:protochlorophyllide reductase (ATP-dependent) iron-sulfur ATP-binding protein | 95.65% |
| magnesium protoporphyrin IX methyltransferase | 95.65% |
| BCD family MFS transporter | 95.65% |
| photosynthetic reaction center subunit H | 69.57% |
| photosynthetic complex assembly protein | 60.87% |
| magnesium-protoporphyrin IX monomethyl ester (oxidative) cyclase | 95.65% |
| DUF3623 domain-containing protein | 100.00% |

**Supplementary Table 3.** Assembly statistics for *Altererythrobacter* sp. DSM24483. Statistics calculated using QUAST.

| assembly | total length | # of contigs | Largest contig | N50 | GC% |
| --- | --- | --- | --- | --- | --- |
| ABySS | 3,583,156 | 523 | 375472 | 177,281 | 62.96 |
| Velvet | 4,549,930 | 33942 | 843 | 551 | 62.95 |
| SPAdes (short read only) | 3,505,755 | 126 | 818185 | 225,709 | 62.96 |
| SPAdes (hybrid assembly) | 3,551,944 | 70 | 2621067 | 2,621,067 | 62.93 |
| Unicycler | 3,482,249 | 26 | 1312088 | 477,084 | 62.97 |
| Flye/Polypolish | **3,668,998** | **8** | **3429213** | **3,429,213** | **60.84** |

**Supplementary Table 4.** Assembly statistics for *Erythrobacter* sp. DSM25594. Statistics calculated using QUAST

| assembly | total length | # of contigs | Largest contig | n50 | GC% |
| --- | --- | --- | --- | --- | --- |
| Abyss | 3,555,036 | 314 | 270603 | 154,259 | 60.78 |
| Velvet | 3,161,924 | 24155 | 553 | 525 | 65.07 |
| SPAdes (short read only) | 3,505,103 | 138 | 281976 | 158,763 | 60.76 |
| SPAdes (hybrid assembly) | 3,551,223 | 57 | 3426610 | 3,426,610 | 60.78 |
| Unicycler | **3,551,136** | **4** | **3428904** | **3,428,904** | **60.79** |

**Supplementary Figure 8.** Gene order plot for regions flanking the photosynthetic gene cluster in *Aurantiacibacter marinus*. Photosynthetic gene cluster is indicated in orange.

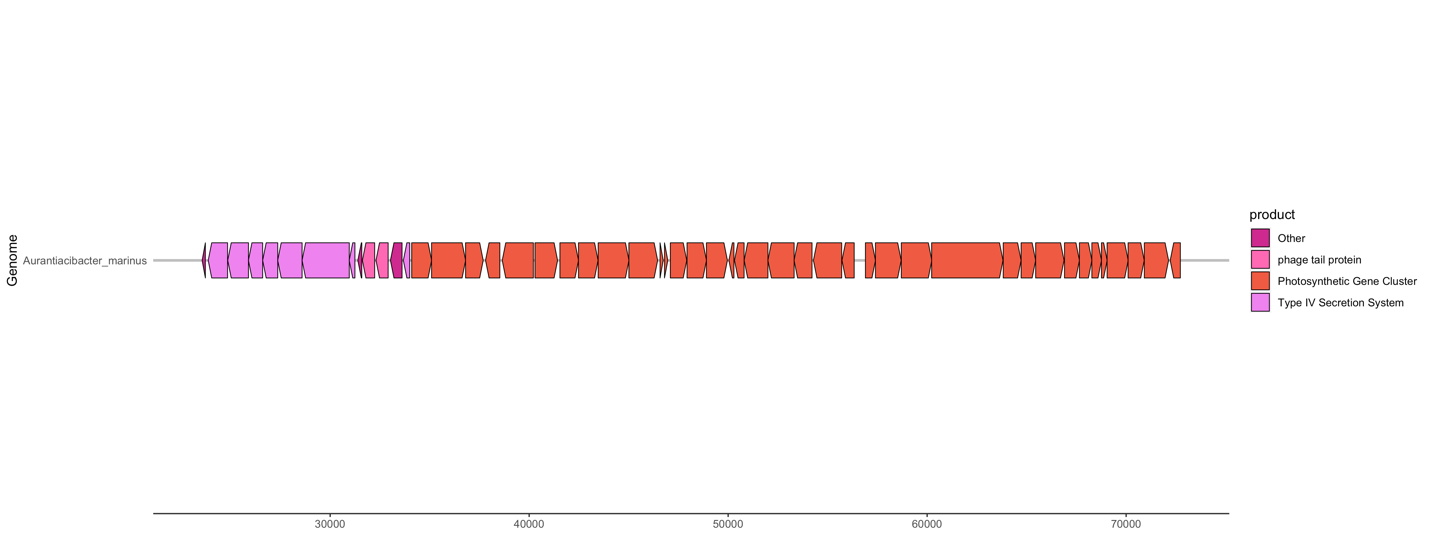

**Supplementary Figure 9.** Pairwise genome alignment for regions flanking the photosynthetic gene cluster in *Aurantiacibacter odishensis* and *Aurantiacibacter zhengii*. Photosynthetic gene cluster is indicated in orange.

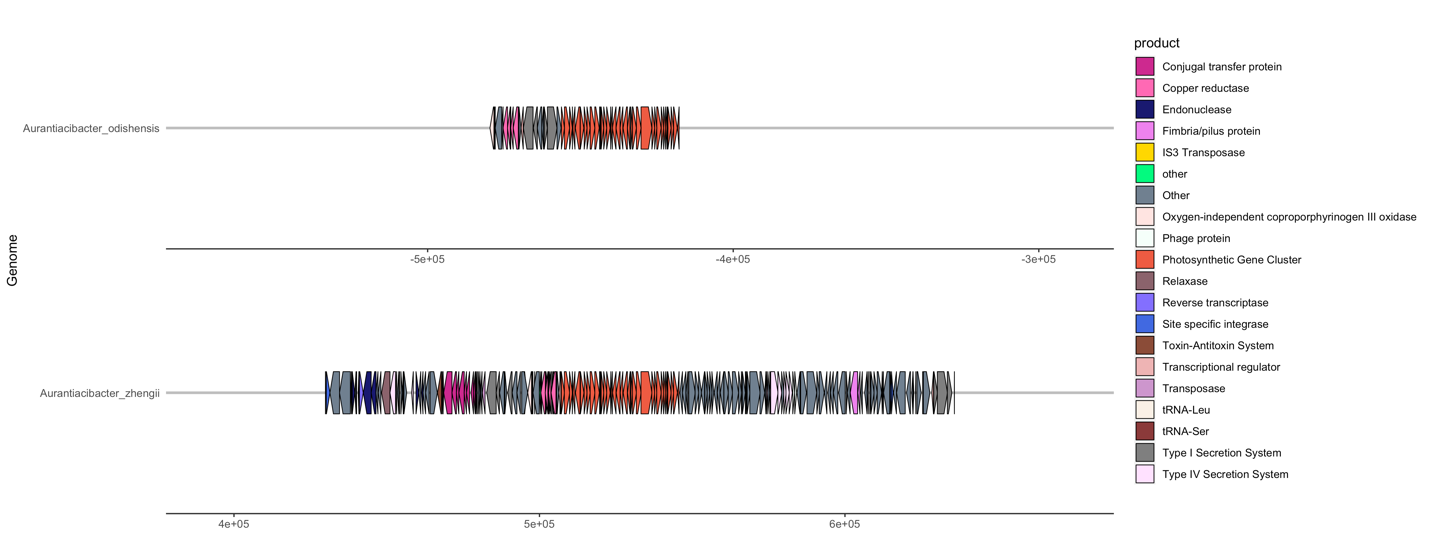

**Supplementary Figure 10.** Gene order plot for *Aurantiacibacter spongiae*. Photosynthetic gene cluster indicated in orange.

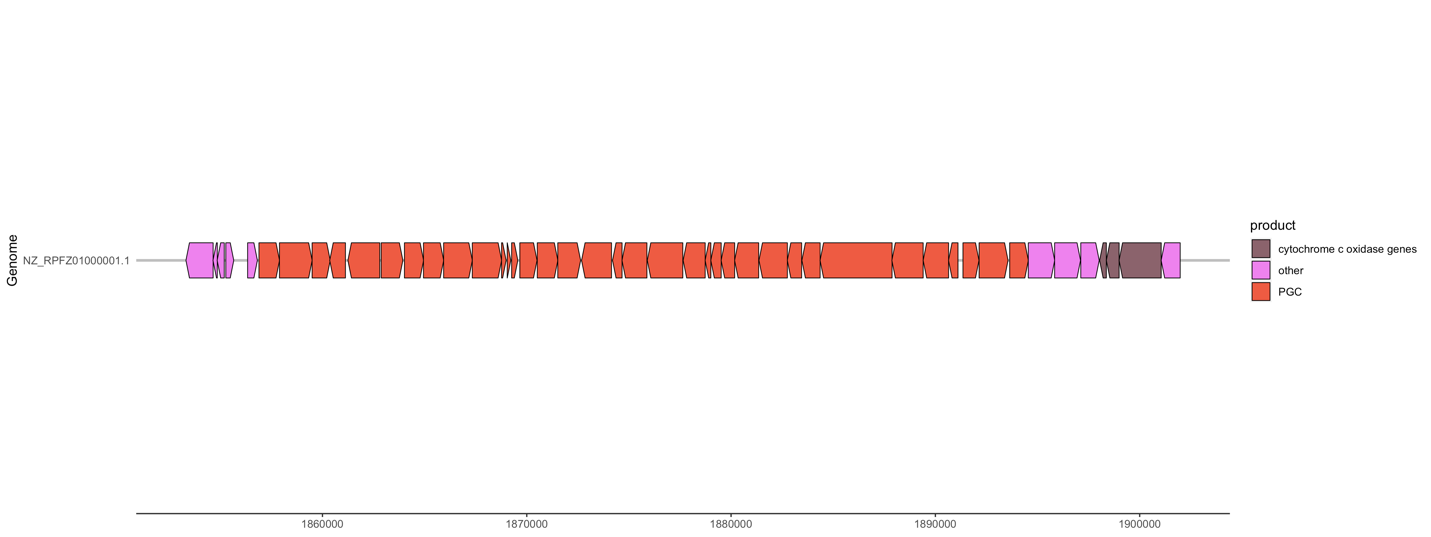

**Supplementary Figure 11.** Gene order plot for *Pontixanthobacter aestiaquae*. Photosynthetic gene cluster indicated in orange.

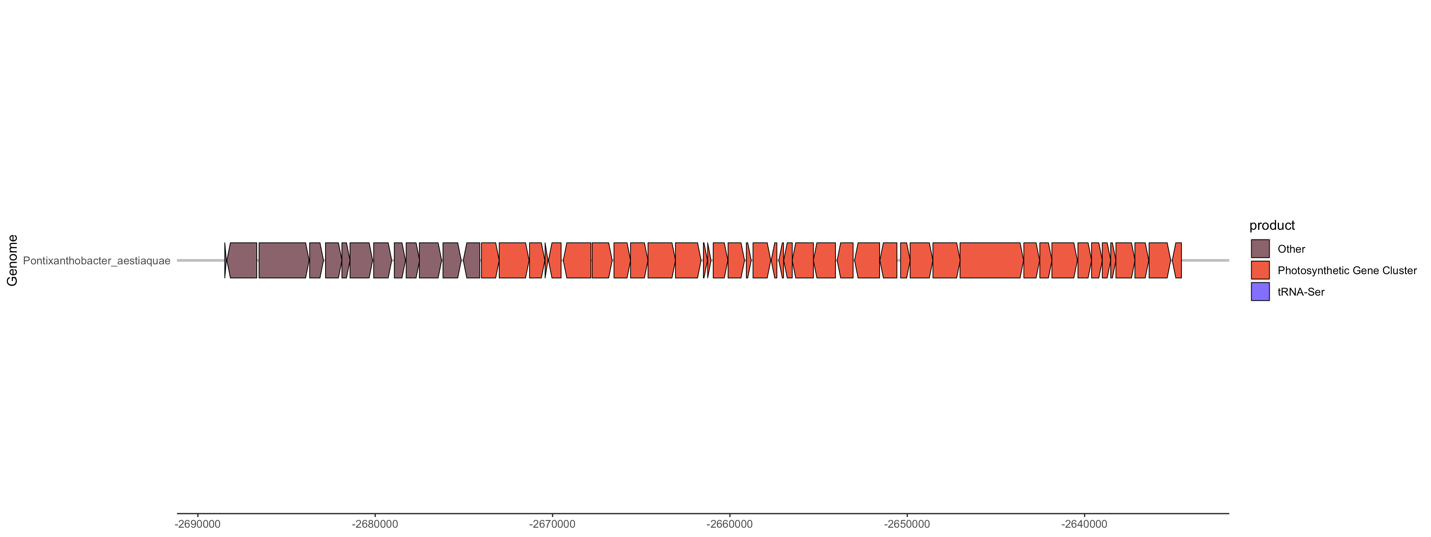

**Supplementary Figure 12.** Gene order plot for *Pontixanthobacter sediminis.* Photosynthetic gene cluster indicated in orange. The photosynthetic gene cluster has become fragmented and is in two locations on the genome separated by 703037 base pairs.

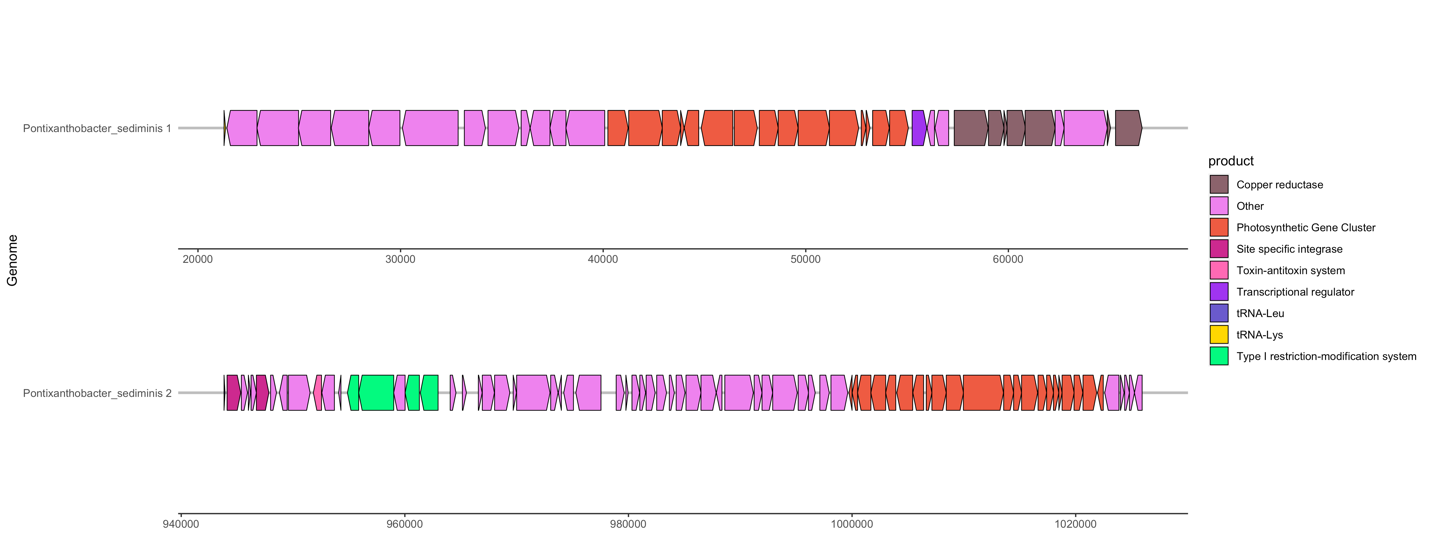
